# Class III PI3K is essential for Wingless secretion and Evi/Wls recycling in *Drosophila*

**DOI:** 10.1101/2025.05.20.655092

**Authors:** Michaela Holzem, Bojana Pavlović, Mareike Munz, Claudia Strein, Jan Gerwin, Marko Lampe, Fillip Port, Michael Boutros

## Abstract

Wnt/Wingless (Wg) signalling is a key regulator of tissue patterning and morphogenesis in *Drosophila*, coordinating cell fate decisions and long-range morphogen signalling. In wing imaginal discs, Wg proteins rely on specialized trafficking machinery, including the cargo receptor Evi/Wls, and are secreted via multiple routes, comprising a glypican-dependent Wg pool and an endocytosis-dependent Wg pool. However, the cellular mechanisms controlling Wg secretion and post-endocytic trafficking in *Drosophila* remain incompletely understood. We performed an *in vivo* kinome- and phosphatome-wide CRISPR-Cas9 screen in *Drosophila* wing imaginal discs using endogenous fluorescently tagged Wg as a readout. Genetic perturbations were combined with super-resolution microscopy and *ex vivo* pharmacological treatments to resolve Wg and Evi/Wls secretion dynamics. We identified Vps15, a core subunit of the class III phosphatidylinositol 3-kinase (PI3K (III)) complex, as a critical factor of Wg secretion. Loss of Vps15 caused apical accumulation of Wg in Wg-secreting cells, selectively impairing an endocytosis-dependent Wg pool while leaving a glypican-mediated Wg pool intact. In contrast to Wg, Evi/Wls did not accumulate in PI3K (III) mutant cells, but instead was subject to proteasome-dependent degradation. Super-resolution imaging further revealed frequent spatial separation of Wg and Evi/Wls in Wg-secreting cells prior to the uptake of the endocytosis-dependent Wg pool. Our study establishes PI3K (III) perturbation as a powerful approach to dissect distinct Wg secretion routes in *Drosophila* wing imaginal discs. We uncovered divergent post-endocytic fates of Wg and Evi/Wls upon perturbation and provided new mechanistic insight into Wg-Evi/Wls dynamics.

## BACKGROUND

The Wnt signalling pathway is one of the key evolutionary innovations at the origin of metazoans (Holstein, 2012; Holzem et al., 2024; Loh et al., 2016). Wnt signalling is essential for the patterning the body axes of all multicellular animals and is involved in cell cycle regulation, genome integrity, cell polarity, organismal development and adult homeostasis (Hayat et al., 2022; Logan and Nusse, 2004). Wnt signalling can be divided into three evolutionarily conserved modules: (i) the secretion of Wnt ligands, (ii) the trafficking of Wnt molecules to receiving cells, and (iii) the reception and activation of intracellular signalling cascades in receiving cells (Alvarez-Rodrigo et al., 2023; Mittermeier and Virshup, 2022). Wnt ligands have been demonstrated to have autocrine, paracrine and endocrine effects (Clevers and Nusse, 2012; Wiese et al., 2018). Upon binding to receptors and co-receptors, Wnt ligands activate different downstream signalling cascades, such as canonical or non-canonical Wnt pathways. Diverse Wnt ligands in combination with various receptors lead to different signalling outcomes, enabling Wnt signalling pathways to activate different physiological programs in a context-dependent manner (Niehrs, 2012; Rim et al., 2022; Wiese et al., 2018).

Wnt/Wg secretion initiates in the endoplasmic reticulum (ER) where Wingless (Wg, human Wnt1) is palmitoylated by the O-acetyltransferase Porcupine (Porcn) and loaded onto the cargo-receptor Evenness interrupted/Wntless (Evi/Wls) (Bänziger et al., 2006; Bartscherer et al., 2006; Goodman et al., 2006; Kadowaki et al., 1996). The Evi/Wls-Wg complex is shuttled via COPII vesicles to the Golgi apparatus and further towards the apical plasma membrane. These vesicles fuse with the plasma membrane and expose Wg to the extracellular matrix (ECM). On the apical membrane, Wg protein can traffic directly via e.g. glypicans such as Dally and Dally-like (Dlp) towards receiving cells (Grobe and Guerrero, 2020; Han et al., 2005; Khare and Baumgartner, 2000), or can be re-internalized and shuttled via the endosomal pathway into multivesicular bodies (MVBs), which can lead to the apical or basal secretion of Wg on exosomes (Beckett et al., 2013; Gross, 2021; Gross et al., 2012). Endocytosis of Wg at the apical membrane is the critical step for its entry into the endosomal pathway and is dynamin-clathrin dependent specifically in Wg-secreting cells (Brunt and Scholpp, 2018; Pfeiffer et al., 2002; Strigini and Cohen, 2000). Further, secretion on specialized cellular protrusions, such as cytonemes was also observed in other model organisms (Routledge and Scholpp, 2019; Stanganello et al., 2015). Upon binding of Wg to Frizzled (Fz) receptors and LRP5/6 co-receptors (Arrow in *Drosophila*) on the receiving cell, intracellular Dishevelled (Dsh) proteins are recruited to the receptor complex, activating the assembly of the so-called signalosome (consisting of CK1a, APC, Shaggy (Sgg, GSK3β in vertebrates), Axin and Dsh), which prevents the degradation of Armadillo (Arm, beta-catenin in vertebrates). Arm can subsequently translocate into the nucleus and activate transcription of target genes such as *distalless, senseless, vestigial* or *naked* and others in a concentration and cell-type dependent manner (Alvarez-Rodrigo et al., 2023; Mittermeier and Virshup, 2022; Swarup and Verheyen, 2012; Widmann and Dahmann, 2009).

Since the discovery of the pathway in the 1980s, many components and distinct mechanisms have been discovered (Baker, 2024; Jenny and Basler, 2014) and a complex picture of Wnt secretion and signalling has emerged. Genetic screens have identified important components of the Wnt signalling pathway in *Drosophila*. In the early days, forward genetic screens were used to identify the underlying genes for specific phenotypes by broad genome mutagenesis (Nüsslein-Volhard and Wieschaus, 1980; Nüsslein-Volhard et al., 1984). Later, screens were based on RNA interference (RNAi) and systematically targeted specific subsets of genes, such as kinases, phosphatases or transcription factors, resulting in the identification of additional Wnt pathway components (Chaudhary and Boutros, 2019; Gross et al., 2012; Swarup et al., 2015). However, these approaches are limited to genes for which the partial knockdown typically induced by RNAi is sufficient to produce detectable phenotypes. Approaches using CRISPR/Cas9 might overcome these limitations by fully knocking out a gene of interest and allow analysis of its cellular mechanisms (Evron et al., 2021; Port and Boutros, 2022). Still, various questions remain unresolved in *Drosophila* Wnt signalling mechanisms and function. For example, the precise mechanism of Wg secretion at the DV boundary, including its loading onto the glypicans Dally or Dlp and the mechanism of endocytosis remain incompletely characterized. Further, the spatial and temporal separation between Wg and the cargo-receptor Evi/Wls also remains controversial.

Here, we identify novel components involved in Wnt secretion in the polarized cells of the *Drosophila* wing imaginal disc by CRISPR mutagenesis screening. We performed a kinome and phosphatome-wide Cas9-mediated mutagenesis screen to identify proteins which influence the secretion of Wg from the epithelium. Our screen identified Vps15 (CG9746) as a gene required for correct protein trafficking of Wg.

Vps15 (previously known as *ird1* in *Drosophila* and PIK3R4 in humans) is the regulatory subunit of the class III phosphatidylinositol 3-kinase (PI3K (III)), also known as Vps34-complex (Panaretou et al., 1997; Shravage et al., 2013; Wu et al., 2007). This complex regulates multiple vesicle-dependent transport processes within the cell, including endocytosis, endosomal trafficking, and autophagy (Abe et al., 2009; Juhász et al., 2008; Lindmo et al., 2008; Ratliff et al., 2015). Class III PI3K phosphorylates phosphatidylinositol (PI) to generate phosphatidylinositol 3-phosphate (PI3P), a lipid that both facilitates vesicle maturation through its membrane localisation (He et al., 2017; Posor et al., 2022) and serves as a binding platform for FYVE- or PX-domain containing effector proteins (Chandra et al., 2019; Hayakawa et al., 2004; Hurley, 2006; Lemmon, 2003). Proteins that bind membranes e.g. via FYVE domains, such as Rabenosyn-5 (Rbsn-5; also known as EEA1) (Abe et al., 2009; Mills et al., 1998; Patki et al., 1997; Simonsen et al., 1998; Stenmark et al., 1996), often function as adaptor proteins that recruit additional binding partners, including Rab5 (Backer, 2008; Nielsen et al., 2000) or SNARE proteins (Backer, 2008; Linnemannstöns et al., 2020).

Here, we sought to determine how PI3K (III) regulates Wg secretion at the DV boundary of the *Drosophila* wing imaginal disc and whether it differentially controls distinct Wg trafficking routes. We hypothesized that PI3K III specifically promotes the maturation of Wg-loaded endocytic vesicles in Wg-secreting cells. By combining *in vivo* CRISPR mutagenesis with super-resolution imaging, we identify PI3K (III) as a key regulator of endocytosis-dependent Wg trafficking. Furthermore, we reveal distinct effects on the cargo receptor Evi/Wls, whose post-endocytic dynamics diverge from those of Wg upon Vps15 perturbation. Together, our findings provide new mechanistic insight into the regulation of Wg secretion and the differential trafficking behaviour of Wg and Evi/Wls in Wg-secreting cells.

## RESULTS

### A CRISPR-Cas9 screen to identify novel genes for Wnt secretion in *Drosophila*

To identify novel genes involved in Wg dynamics, we performed an *in vivo* genetic screen in *Drosophila* imaginal wing discs using tissue-specific CRISPR-Cas9 mutagenesis (Port et al., 2020). We combined an optimized Cas9 transgene with sgRNAs of the HD_CFD library, a large-scale collection of sgRNA transgenes, each expressing two sgRNAs targeting two independent positions per gene (Port et al., 2014; Port et al., 2020). Cas9 expression was induced by the *hedgehog-Gal4 (hhGal4)* driver specifically in the posterior half of the wing disc, which allows for the comparison of perturbed and unperturbed cells side-by-side within the same tissue (Fig. 1a). Acute CRISPR mutagenesis in somatic tissues leads to the generation of small insertions and deletions (indels) at the target locus, which frequently induces loss-of-function alleles. However, the heterogeneous nature of the induced mutations and limitations of CRISPR activity typically result in genetic mosaics, where cells with and without gene disruption are present in the same tissue (Port and Boutros, 2022). To visualize Wg protein, we utilized a previously described insertion of GFP in the endogenous *wg* locus (Wg::GFP) (Port et al., 2014). In addition, we used a transcriptional reporter of the Wnt target gene *distalless* (*dll*, Dll::dTom) (Chaudhary et al., 2019) (Fig. 1a). In control cells, which expressed sgRNAs targeting a gene not expressed in wing imaginal discs, Wg::GFP and Dll::dTom were present at similar levels in the anterior and posterior compartment of the wing disc (Fig. 1b-b’’). In contrast, cells which expressed Cas9 and *Evi/Wls-sgRNAs* had strongly elevated levels of Wg::GFP in Wnt producing cells, a consequence of a block in protein secretion (Bänziger et al., 2006; Bartscherer et al., 2006), and showed reduced expression of the Wnt target gene *dll* (Fig. 1c-c’’).

**Figure 1.**
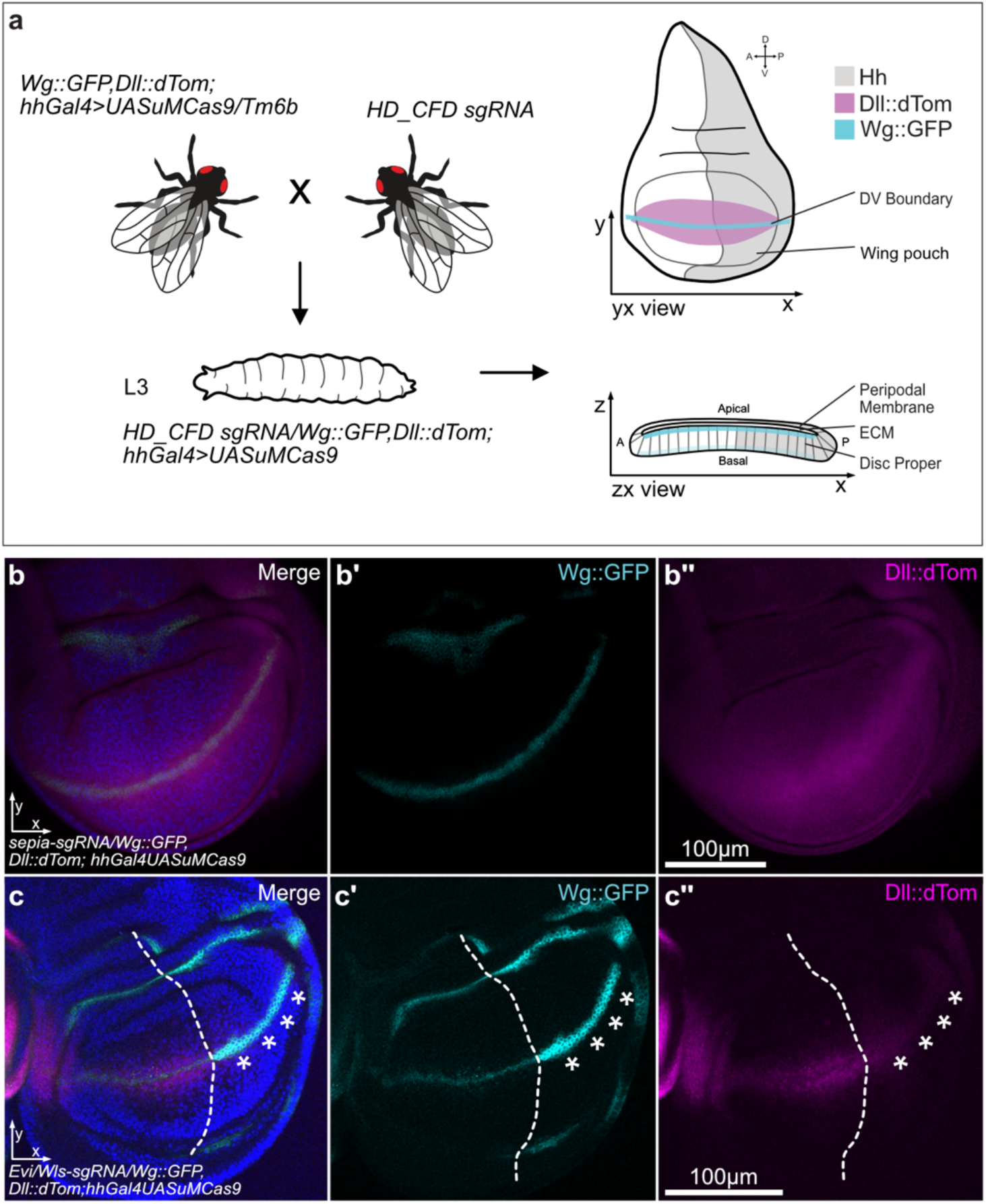
Screening for novel candidates of Wg secretion at the *Drosophila* DV boundary. **(a)** Overview of the screening procedure. The driver line *w-;Wg::GFP,Dll::dTom;hh-Gal4,UAS-Cas9uM/TM6b* was crossed to the sgRNA containing HD_CFD fly lines. Third instar larvae (wandering stage) were dissected and wing imaginal discs were mounted for confocal analysis using the endogenously tagged genes *wg* and *dll* as readout. Indicated are also the orientation and morphology of the wing imaginal disc in the YX view which will be used throughout the manuscript. Additionally, we indicated the ZX view which we use to show protein localisation at the DV boundary and through the disc proper. Expression patterns of Wg::GFP (cyan), Dll::dTom (magenta) and hh (grey) are indicated schematically. **(b-b’’)** Confocal imaging of the negative control *sepia-sgRNA* (n=9) with the endogenous labelled **(b’)** Wg::GFP (cyan) and **(b’’)** Dll::dTom (magenta) pattern. No differences in fluorescence signal are detected in the anterior or posterior compartment. **(c-c’’)** Confocal imaging of the positive control line *Evi/Wls-sgRNA* (n=4) showing a strong increase of **(c’)** Wg::GFP (cyan) at the DV boundary (asterisks), and **(c’’)** a loss of Dll::dTom (magenta) in the corresponding area. The dashed line indicates the border between the anterior and posterior compartment. Presented confocal images are average intensity projections for both genotypes. All nuclei are stained with Hoechst33342 (blue). Confocal images were taken with the 40x oil objective. ECM: Extracellular matrix; DV: Dorso-ventral boundary.

We performed a targeted screen focused on genes encoding kinases or phosphatases, proteins known to be central regulators of intracellular signalling. The HD_CFD library contains sgRNA lines targeting 288 of the 327 kinases and 115 of the 186 phosphatases encoded in the *Drosophila* genome (Suppl. Table S1). These include 51 genes which have been implicated in Wg regulation in previous screens and can be used to estimate the efficiency of the screen (Chaudhary and Boutros, 2019; Gross et al., 2012; Swarup et al., 2015) (Suppl. Table S2).

In total, we screened 605 sgRNA lines targeting 411 unique genes and uncovered 56 genes that reduce or increase Wg in the wing disc. Of these, 50 genes are among the 51 previously described regulators of Wg, representing a 98% overlap and suggesting high efficiency of our CRISPR perturbations. In addition, we identified Wg phenotypes with 6 lines targeting genes so far not implicated in Wg trafficking, which showed an increase or decrease of Wg signal at the DV boundary. The induction of DNA double-strand breaks through CRISPR nucleases is known to result in frequent mitotic recombination (Allen et al., 2021; Brunner et al., 2019), which can result in loss of heterozygosity of the Wg::GFP or Dll:dTom reporters if induced between their locus and the centromere. We therefore rescreened any genes located between the centromere and the *wg::GFP* gene on chromosome arm 2L by *anti-wg* immunohistochemistry. This revealed that the phenotypes for two of the initial hits reflect mitotic recombination rather than genuine regulation of Wg. The remaining four positive hits (Vps15, Mkk4, Takl2, Pink1) showed reproducible effects on Wg protein (Suppl. Table S3).

For further characterization, we focused on Vps15, which exhibited the strongest accumulation of Wg at the DV boundary following CRISPR-mediated mutagenesis. Vps15 is a core subunit of the class III PI3K complex, a key regulator of vesicle-dependent trafficking processes, including endocytosis and autophagy (see Background) (Abe et al., 2009; Juhász et al., 2008; Lindmo et al., 2008; Ratliff et al., 2015).

### Wg protein accumulates at the apical surface in Vps15 mutant wing imaginal discs

Strong accumulation of Wg was observed at the DV boundary and in additional Wg expression domains within Vps15-perturbed posterior wing disc tissue using HD_CFD Vps15-sgRNA#009 (n=13; Fig. 2a,a′). To validate this phenotype, we analysed a second sgRNA line from an independent sgRNA library (Vps15-sgRNA#83822; BDSC), which produced a comparable phenotype (n=13; Fig. 2c,c′). In both cases, Wg accumulation was not uniform across the posterior compartment, likely reflecting genetic mosaicism resulting from acute CRISPR-mediated mutagenesis. Further analysis of Wg subcellular localization in Vps15-perturbed cells revealed that Wg predominantly accumulates in large punctate structures at or near the apical membrane (Fig. 2b,b′ and d,d′).

**Figure 2.**
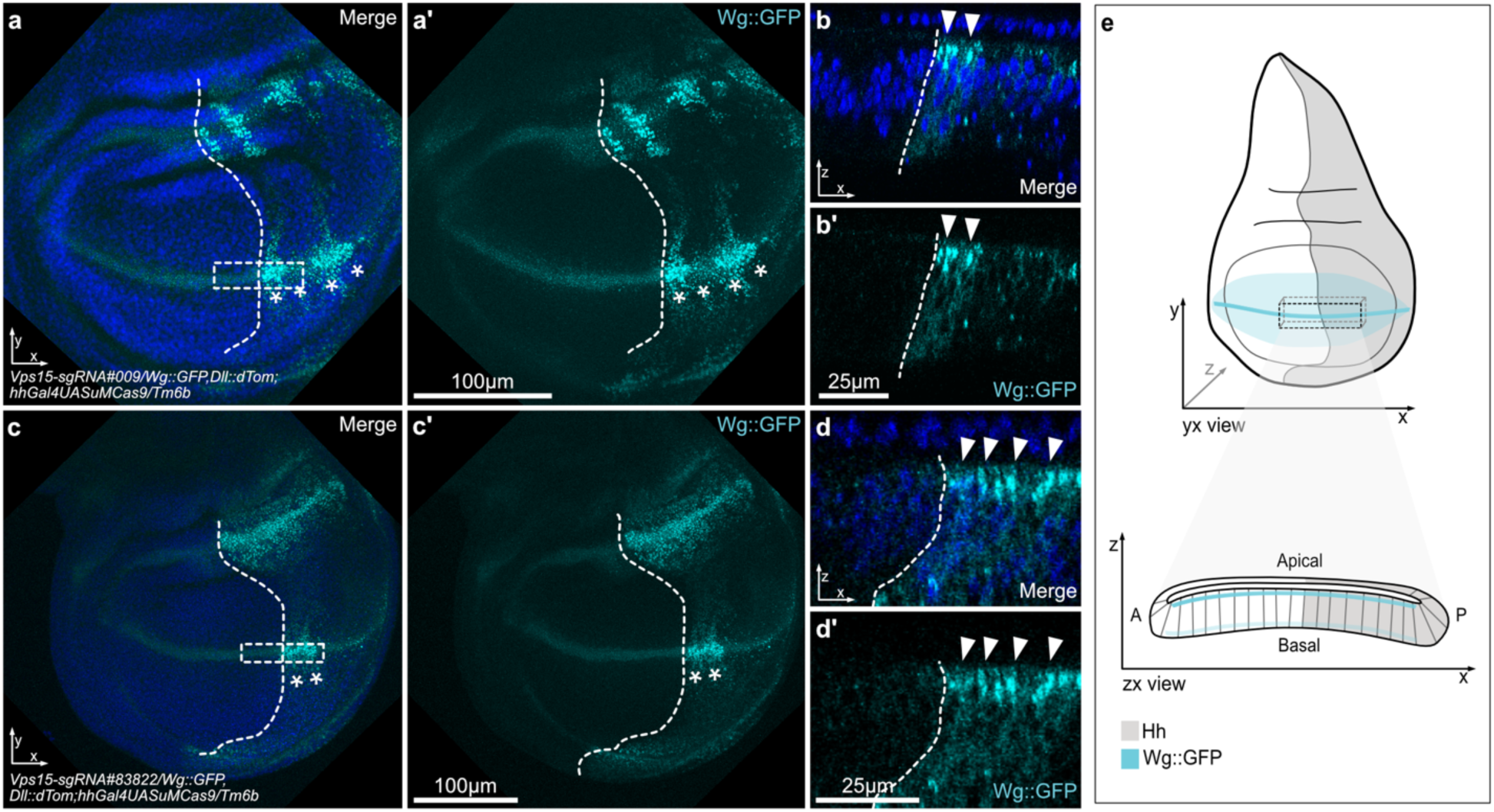
Characterisation of Vps15 mutagenesis in *Drosophila*. **(a,a’)** Wg protein (cyan) abundance at the DV boundary in the wing discs of larvae of the cross *w-;Wg::GFP,Dll:dTom;hh-Gal4,UASCas9uM/Tm6B* with the HD_CFD *Vps15-sgRNA#009* fly line. The perturbed posterior area of the wing disc is indicated by the dashed line. Genetic mosaics with Vps15 mutagenesis are labelled by white asterisks. **(b,b’)** The Wg protein (cyan) accumulation in the ZX view shows its location in the apical area of the cells (arrowheads). **(c,c’)** Wg phenotype (cyan) of the BDSC *Vps15-sgRNA#83822* fly line crossed with *w-;Wg::GFP,Dll:dTom;hh-Gal4,UASCas9uM/Tm6B*. Strong accumulation can also be detected in Wg-secreting cells at the DV boundary (asterisks) and **(d,d’)** in the apical membrane area in the ZX view (arrowheads). **(e)** Schematic overview of wing imaginal disc orientation. The full wing imaginal disc is shown with the orientation used in all presented images. Wg expression at the DV boundary is indicated in cyan, along with the Wg morphogen gradient (semi-transparent cyan). Below, a YX view of the wing disc at the DV boundary is shown which corresponds to the boxed area of the wing imaginal disc. In the corresponding ZX view, only Wg-secreting cells (cyan) are visible. **(a-d)** All confocal images were taken with the 63x oil objective. Nuclei were stained with Hoechst33342 (blue). All discs are orientated with the wildtype tissue towards the left (anterior) and perturbed tissue towards the right (posterior). Confocal images are maximum intensity projections for both genotypes in the YX view and single slice images in the ZX view.

We further confirmed effective perturbation of Vps15 for both sgRNAs by Sanger sequencing and analysed the results for their knockout efficiency. On average, 40.9% (±7.23%) of sgRNA#009 and 40.3% (±4.88%) of sgRNA#83822 samples showed deletions at or near the respective sgRNA cut sites (see Suppl. Table S4). Both sgRNAs efficiently induced mutagenesis of Vps15; however, sgRNA#83822 exhibited greater consistency, as indicated by its lower standard deviation. Based on these sequencing results and our additional confocal analyses (Fig. 2), we therefore primarily used the sgRNA#83822 line throughout the manuscript for Vps15 mutagenesis, as it consistently produced a strong and reliable perturbation of Wg in Wg-secreting cells.

To determine whether Wg accumulation in Wg-secreting cells at the DV boundary results from increased gene expression, we performed fluorescent *in situ* hybridization (FISH) to analyse *wg* transcript levels. No increase in *wg* mRNA was detected, indicating that the elevated Wg protein levels arise from defects in protein trafficking, increased translation, or reduced protein turnover, rather than transcriptional upregulation (Suppl. Fig. S1a,b). We next assessed whether RNAi-mediated depletion of Vps15 could phenocopy the CRISPR-induced Wg accumulation, using the same *hh*-*Gal4* driver employed in the CRISPR screen. None of the three publicly available Vps15 RNAi lines tested produced a Wg phenotype comparable to that observed with either sgRNA line (Suppl. Fig. S1c-e). These results suggest that RNAi-mediated knockdown alone is insufficient to reduce Vps15 levels below the threshold required to disrupt Wg trafficking. Consistent with this interpretation, previous studies investigating RNAi-mediated depletion of PI3K (III) associated components such as Atg6 (Lőrincz et al., 2014) required co-expression of Dice, which can increase RNAi efficiency.

To verify functional disruption of PI3K(III), we used a UAS-2xFYVE-GFP reporter that labels PI3P-positive endocytic vesicles via the FYVE domain. Class III PI3K generates PI3P, a lipid required for vesicle maturation and recruitment of FYVE-domain proteins (Chandra et al., 2019; Hayakawa et al., 2004; Hurley, 2006; Lemmon, 2003). We introduced this reporter into our CRISPR/Cas9 system and compared GFP fluorescence between control sepia perturbation and Vps15#83822-perturbed tissue. GFP signal was lost upon Vps15 perturbation, confirming functional impairment of PI3K(III) activity in our system (Suppl. Fig. S2).

### Effects of Vps15 loss on adult wing morphology

Perturbations of Wnt signalling and loss of Wg function are known to affect morphogenesis of the adult wing. These phenotypes can range from a reduction of wing margin bristles, to notches of the margin and up to the complete loss of adult wings. Wings can therefore be used as a phenotypic readout for malfunctioning Wnt signalling. Mutagenesis of Vps15 is lethal during late larval/pupal stages when induced by the *hh-Gal4* driver, precluding analysis of adult wings. We therefore induced CRISPR mutagenesis with a different Gal4 driver, with an expression pattern more restricted to the developing wing (*w-;; pdm2-Gal4,UASCas9uM/Tm6B*). Crosses with the *Vps15-sgRNA#009* or *Vps15-sgRNA#83822* lines resulted in viable flies with malformed wings. For *Vps15-sgRNA#009,* we observed a vein phenotype in 64.9% of females (n=23) and 57.7% of the male offsprings (n=12). In male wings specifically, we also detected bubbles (15,4%; n=4), where the attachment between the dorsal and ventral layer of the wing is disrupted, as well as some notches (3.8%; n=1) (Suppl. Fig. S3a,b). No disturbed bristles patterns were observed in these samples. 35.1% (n=13) of female and 23.1% (n=6) of male wings did not show a phenotype and resembled wildtype wings (Suppl. Fig. S3d). For *Vps15-sgRNA#83822* line, 100% (n=32 for both sexes) of all female and male wings showed a phenotype. All wings were malformed and also showed a vein phenotype. Additionally, 15.6% (n=5) of female and 28.1% (n=9) of male wings showed notches at the wing margins, including a loss of bristles in the corresponding area (Suppl. Fig. S3c,d). A disturbed bristle pattern was detected in 28.1% (n=9) of female and 31.3% (n=10) of male wings, without forming a notch (Suppl. Fig. S3d). Overall, perturbation of Vps15 with either sgRNA line resulted in wing abnormalities, some of which are consistent with a disruption of Wnt signalling. While some phenotypes are consistent with reduced Wg signalling, overall, the morphological changes, such as deregulation of wing vein formation, indicate that also other processes are perturbed.

### Loss of PI3K (III) subunits drives apical Wg accumulation

The class III PI3K complex includes Vps15, Atg6, and the catalytic subunit Vps34 (see Background section). To determine whether the observed Wg phenotype following Vps15 mutagenesis reflects a loss of PI3K (III) function, we performed CRISPR mutagenesis using sgRNAs targeting Atg6 (Atg6-sgRNA#80830, BDSC) and Vps34 (Vps34-sgRNA#92484, BDSC). Consistent with previous reports (Abe et al., 2009; Lőrincz et al., 2014), loss of either subunit caused pronounced Wg accumulation in Wg-secreting cells at the DV boundary, with Wg predominantly localized apically, closely phenocopying the effects observed upon Vps15 perturbation (Suppl. Fig. S4). Together, these findings indicate that proper Wg trafficking depends on the coordinated activity of all PI3K (III) subunits, as perturbation of Vps15, Atg6, or Vps34 equally disrupts Wg localization in Wg-secreting cells in the wing imaginal disc.

### Vps15 mutagenesis alters extracellular Wg dynamics in the wing imaginal disc

Class III PI3K has been shown to be essential for endocytic vesicle maturation, as previously described in other tissues (Abe et al., 2009; Juhász et al., 2008; Lindmo and Stenmark, 2006; Shravage et al., 2013). Secreted Wg forms a long-range pool delivered via glypicans (will be referred to as “direct” Wg pool) and a short-range pool that is endocytosed by the secreting cells and trafficked through endosomes (will be referred to as “indirect” Wg pool). We hypothesized that PI3K (III) perturbation would specifically disrupt the apically endocytosed “indirect” pool, while leaving the apical glypican-mediated “direct” pool largely intact. To test this, we first examined whether the apical Wg gradient formation, which is maintained by the “direct” pool, is disrupted upon Vps15 mutagenesis. We employed an immunohistochemistry protocol without permeabilization to selectively label extracellular Wg (Wg^Ex^), thereby excluding intracellular signal (Strigini and Cohen, 1999). In parallel, the endogenous tag of Wg::GFP allowed visualization of total Wg protein and identification of Vps15-perturbed cells within the same tissue. Wg^Ex^ was detected in both wildtype and Vps15 perturbed tissue, with stronger Wg^Ex^ signal observed in the perturbed posterior compartment (Fig. 3a-a’’, n=5). Fluorescence intensity measurement confirmed the significant increase of Wg^Ex^ in the Vps15 perturbed tissue (Fig. 3b, n=5 and Suppl. Fig. S5a). Analysis of YX projections revealed that Wg^Ex^ localized slightly apical to the bulk of total Wg::GFP (stained with anti-GFP), consistent with its extracellular distribution (Fig. 3c-c’’, n=5). These results indicate that loss of Vps15 does not impair the formation or maintenance of the “direct” Wg pool.

**Figure 3.**
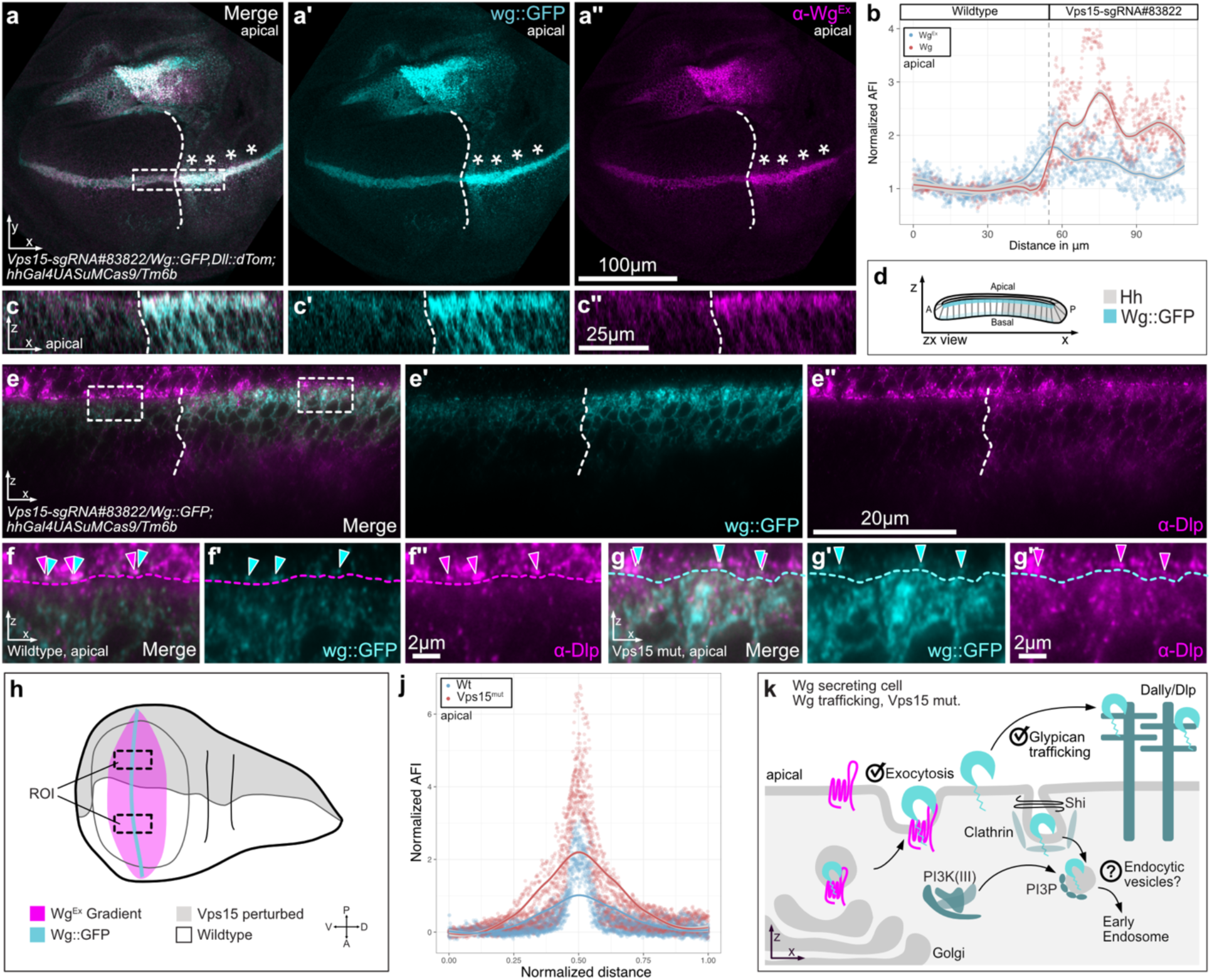
Effects of Vps15 mutagenesis on extracellular Wg and gradient formation. **(a-a’’)** Extracellular Wg (Wg^Ex^, magenta) accumulates at the DV boundary in Vps15 mutated clones (asterisks, n=5; *w-;Vps15-sgRNA#83822/Wg::GFP,Dll::dTom;hhGal4UASuMCas9/Tm6b*). **(b)** Average fluorescence intensity is significantly increased for both total Wg::GFP (red, p<0.0001) and extracellular Wg^Ex^ (blue, p<0.0001) in Vps15 perturbed regions, although Wg^Ex^ signal remains lower than total Wg staining (n=5). For statistical test please refer to Suppl. Fig. S5. **(c-c’’)** The ZX view revealed apical accumulation of Wg^Ex^ in Vps15 mutant cells, positioned above the endogenous Wg::GFP signal (white arrowheads, n=5). **(d)** Schematic orientation of the ZX view as reference for this figure. **(e-e’’)** Immunofluorescence of Dlp (magenta) and Wg::GFP (cyan) in the *w-;Vps15-sgRNA#83822/Wg::GFP; hhGal4UASuMCas9/Tm6b* indicates transfer of Wg onto glypicans using STED super-resolution microscopy (n=6). **(f-f’’)** Higher-magnification view of the wildtype apical region. The apical membrane is inferred from Dlp staining (magenta dashed line), and close spatial proximity between Dlp (magenta arrowheads) and Wg (cyan arrowheads) is observed. **(g-g″)** High-magnification view of the Vps15 perturbed region also shows close proximity between Dlp (magenta arrowheads) and Wg (cyan arrowheads) above the apical membrane, inferred from Wg staining and indicated by a cyan dashed line. **(h)** Schematic illustration of the gradient measurement strategy. Identical regions of interest (ROIs) were analysed in unperturbed and Vps15 perturbed tissue, and Wg^Ex^ fluorescence intensity was used to assess gradient range. **(j)** Normalized average fluorescence intensity measurements of the Wg^Ex^ gradient in unperturbed (blue) and perturbed (red) tissue (n=9). **(k)** Schematic summary of the proposed role of class III PI3K in Wg secretion. Wg is exocytosed into the extracellular matrix, transferred onto glypicans such as Dlp, and forms a morphogen gradient. A potential effect on the endocytosed “indirect” Wg signalling pool remains to be determined (question mark). **(a-f’’)** All pictures are oriented with anterior to the left. The borders of the anterior and posterior area are indicated by a white dashed line. Confocal images are taken with the 63x oil objective. Presented images are maximum intensity projections for the YX view, all ZX views are single slices. Detailed parameters for the super resolution images can be found in Suppl. Table S5. AFI: Average fluorescence intensity; mut: mutated; ROI: Region of interest; Wt: Wildtype.

Consistently, immunofluorescence staining for the glypican Dally-like protein (Dlp), combined with super-resolution microscopy (Fig. 3e-e’’, n=6), revealed close spatial association between Wg and Dlp in both wildtype (Fig. 3f-f’’) and Vps15 perturbed tissue (Fig. 3g-g’’). Together, these findings suggest that Wg exocytosis and its subsequent transfer onto glypicans is likely unaffected by Vps15 loss in Wg-secreting cells (summarized in Fig. 3k).

Finally, we examined the spatial spread of the Wg^Ex^ gradient to assess whether Vps15 perturbation affects the apical long-range Wg trafficking. Wg^Ex^ fluorescence intensity profiles were quantified in both perturbed and unperturbed apical regions of the same imaginal disc (Fig. 3g), with the distribution of intensities reflecting the gradient range. Although the intensity profiles differed in amplitude, consistent with higher Wg^Ex^ levels in the perturbed region, the shape of the apical gradient remained comparable between the regions (Fig. 3h,j; n=9). Thus, the glypican-mediated “direct” Wg pool remains functional, and long-range Wg trafficking is largely unaffected by loss of Vps15 (summarized in Fig. 3k).

### Apical Wg endocytosis occurs independently of class III PI3K activity

The mechanism by which class III PI3K regulates the “indirect” Wg signalling pool in Wg-secreting cells remains unclear. Based on previous studies, we hypothesized that PI3P is required for the maturation of Wg-loaded endocytic vesicles into early endosomes (Fig. 3k, question mark). To test this, we asked whether Wg is still endocytosed in Wg-secreting cells in the absence of Vps15 and whether the apical accumulation of Wg in Vps15-perturbed tissue reflects impaired vesicle maturation. We performed a “Wg uptake assay”, which selectively labels extracellular Wg and detects Wg internalized into secreting cells (Witte et al., 2021). In this assay, endocytosed Wg (Wg^In^) is visible intracellularly, whereas impaired endocytosis results in little or no intracellular signal (Fig. 4a,a’). Using wing discs expressing Wg::GFP, we simultaneously visualized total Wg and Wg^In^ in the same tissue. All imaging was performed with super-resolution microscopy to maximize spatial resolution and allow detailed analysis of Wg dynamics.

**Figure 4.**
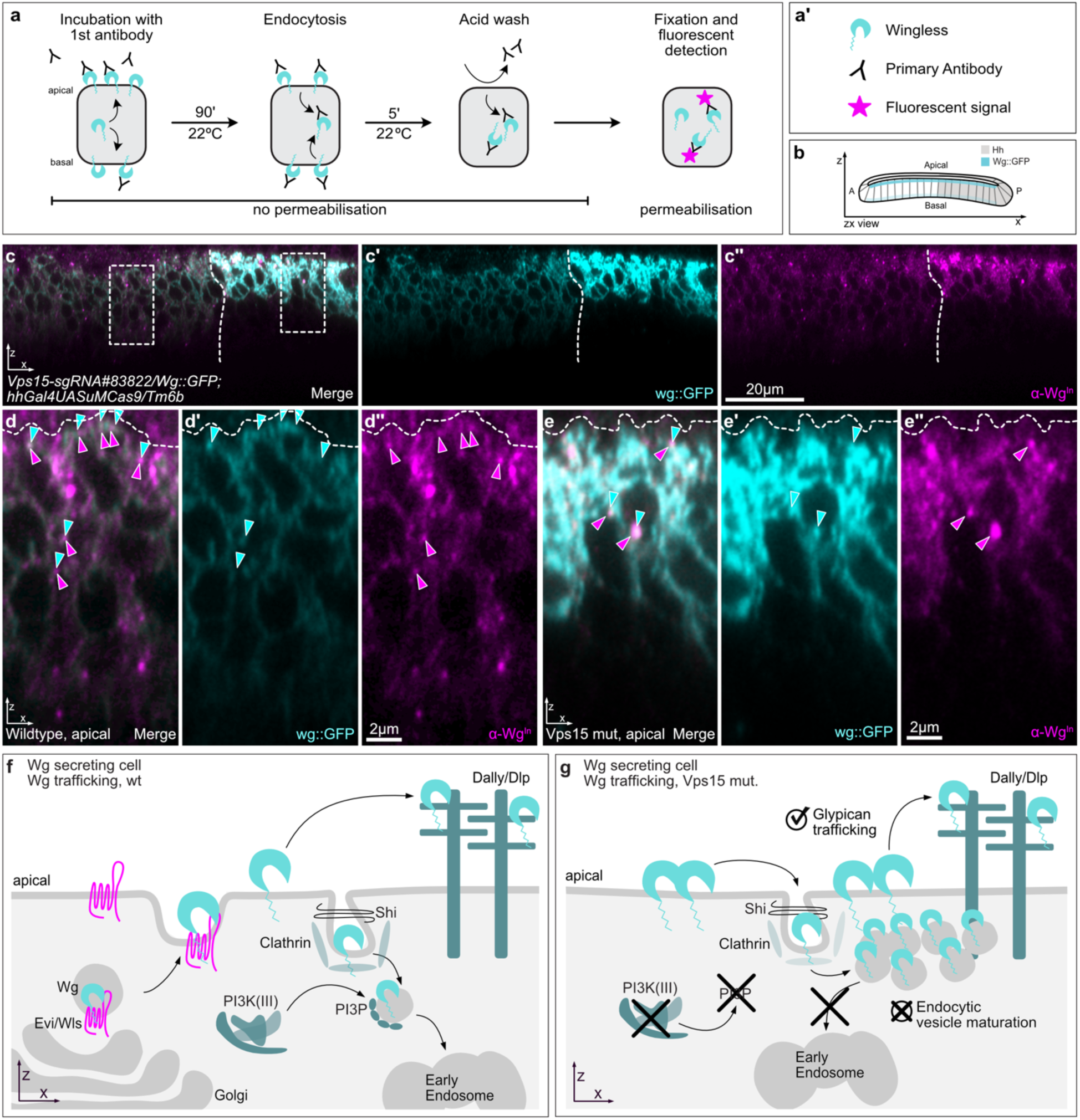
Apical Wg endocytosis in *Drosophila* wing imaginal discs. **(a)** Schematic overview of the “Wg-uptake” assay (Witte et al., 2021). Wing imaginal discs were incubated at 22°C for 90 minutes with a primary antibody to label extracellular Wg and allow its endocytosis. Unbound extracellular primary antibody was subsequently removed by an acid wash. All steps up to this point were performed without permeabilization. The discs were then fixed and incubated with a fluorescently conjugated secondary antibody to detect antibody-labelled Wg. Internalized, antibody-labelled Wg is referred to as Wg^In^. **(a’)** Legend of Wg uptake assay in (a). **(b)** Reference view showing wing disc ZX orientation. **(c-c’’)** Super-resolution images of the ZX view at the DV boundary of treated wing discs, showing the Wg^In^ in Wg-secreting cells (magenta; n=3) in *w-;*Vps15*-sgRNA#83822*/*Wg::GFP;hh-Gal4UASCas9uM/Tm6B*. Unperturbed, anterior tissue is oriented to the left. The hh border between compartments is indicated by a dashed line. Higher-magnification areas are indicated with dashed boxes. **(d-d’’)** Higher-magnification view of wildtype tissue. Total Wg::GFP (cyan arrowheads) is distributed normally and colocalizes with Wg^In^ (magenta arrowheads). Wg^In^ is distributed throughout the disc proper and spreads toward the basal side. **(e-e’’)** Higher-magnification view of Vps15 perturbed tissue. Total Wg::GFP (cyan) accumulates strongly, and Wg^In^ (magenta) clusters primarily in the apical region of the disc proper. Colocalization between total Wg (cyan arrowheads) and Wg^In^ (magenta arrowheads) is observed. These images indicate that Wg endocytosis remains functional in Vps15 perturbed cells, but endocytic vesicles accumulate apically. **(d-e)** Predicted apical membrane is indicated with a white dashed line. **(f)** Schematic overview of wildtype Wg secretion dynamics at the apical membrane. Wg (cyan) is exocytosed and can be transferred to glypicans such as Dally or Dlp. A subset of Wg is also endocytosed from the apical membrane in a dynamin- and clathrin-dependent manner and subsequently enters the endosomal pathway. PI3K (III) phosphorylates PI to PI3P, which is required on endocytic vesicle membranes for maturation into early endosomes. **(g)** Schematic representation of Wg secretion in Vps15-perturbed tissue at the DV boundary. Wg is exocytosed and transferred onto glypicans as in wildtype tissue. Wg is also endocytosed from the apical membrane; however, due to the absence of PI3P on endocytic vesicle membranes, maturation into early endosomes is impaired. As a result, Wg-loaded endocytic vesicles accumulate in the apical region of the disc proper. Detail imaging parameters of the super-resolution images are provided in Suppl. Table S5. mut: mutated; Wt: Wildtype.

In wildtype tissue, Wg^In^-positive puncta were distributed throughout the disc proper of the wing (Fig. 4c–c″; n=3). Since the signal of the total Wg is much weaker than in the perturbed half of the disc, overlapping signals are more difficult to visualize. Still, our analysis shows that mainly all Wg^In^ puncta overlapped with total Wg staining (Fig. 4d-d’’). In the Vps15-perturbed tissue, total Wg::GFP strongly accumulated in Wg-secreting cells at the DV boundary as previously observed. This accumulation is accompanied by pronounced Wg^In^ enrichment in the apical region of the disc proper (Fig. 4e-e’’). In Vps15-mutant tissue, overlap between Wg::GFP and Wg^In^ was also observed. (Fig. 4e). Together, these findings indicate that Wg endocytosis itself is not impaired by loss of class III PI3K activity in Wg-secreting cells. Rather, we hypothesize that endocytic vesicles accumulate apically and fail to mature into early endosomes. Consistent with this idea and previous literature, such vesicle accumulation likely account for the pronounced intracellular Wg signal observed in Vps15-perturbed cells (Fig. 4f,g).

### Vps15 perturbation alters Evi/Wls abundance and trafficking

In addition to Wg, the cargo receptor Evi/Wls is also endocytosed at the apical membrane (Belenkaya et al., 2008; Gasnereau et al., 2011; Port et al., 2008). Whether the “indirect” Wg signalling pool remains associated with Evi/Wls during endocytosis or dissociates after the complex has reached the apical membrane remains controversial (Coombs et al., 2010; Sharma and Chaudhary, 2024). We therefore investigated the secretion and apical trafficking of Evi/Wls in Wg-secreting cells upon Vps15 perturbation. To examine this, we performed immunohistochemistry using a polyclonal anti-Evi/Wls antibody (Port et al., 2008) in Vps15 perturbed tissue expressing endogenously tagged Wg::GFP. In contrast to the strong accumulation of Wg, Evi/Wls did not accumulate in Vps15-mutant tissue (Fig. 5a-a″; n=12). Instead, Evi/Wls protein levels were reduced in the perturbed posterior compartment, despite strong Wg accumulation in the same cells. This reduction was also evident in YX projections, where Evi/Wls levels were decreased in the apical region of the disc proper (Fig. 5b-b″). Fluorescence intensity measurements comparing wildtype and perturbed compartments confirmed a significant reduction of Evi/Wls signal (p<0.0001) upon loss of Vps15 (Fig. 5c and Suppl. Fig. S5; n=12). These results indicate that the effects of Vps15 perturbation on Wg and Evi/Wls are distinct.

**Figure 5.**
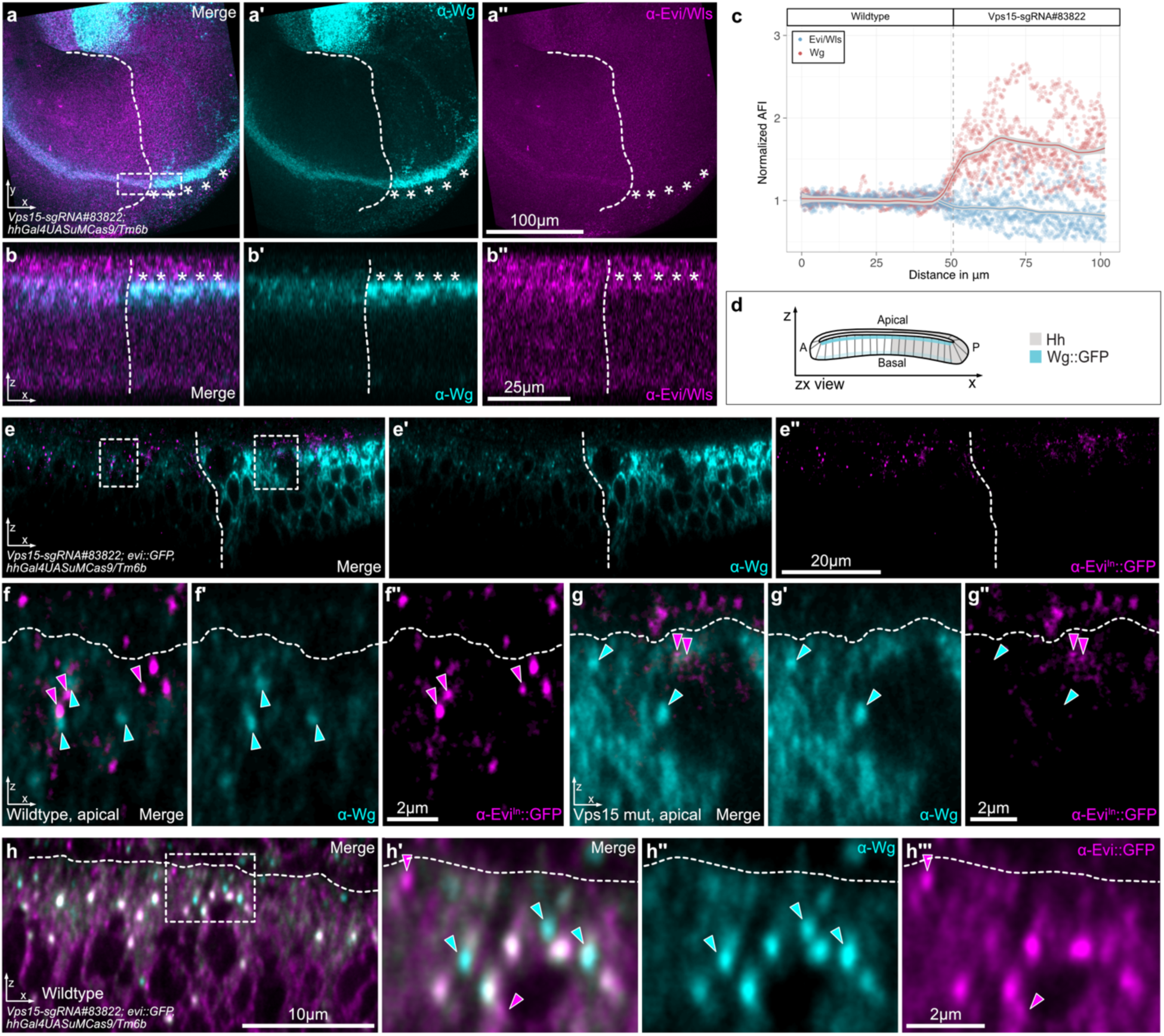
Effects of Vps15 perturbation on Evi/Wls. **(a-a’’)** Evi/Wls expression (magenta) is reduced in Vps15 perturbated cells despite strong accumulation of Wg (cyan, asterisks). Genotype: *w-;Vps15-sgRNA#83822;hhGal4UASuMCas9/Tm6b* (n=12). **(b-b’’)** The Evi/Wls (magenta) reduction is clearly visible in the apical region of the disc proper in the perturbed half of the tissue when viewed in the YX plane. **(c)** Average fluorescence intensity measurements of Wg (red) and Evi/Wls (blue) (n=12) also reveal the reduction of Evi/Wls in the perturbed half of the disc. **(d)** Schematic drawing of the ZX view presented in this figure. **(e-e″)** The Evi/Wls uptake assay was visualized using super-resolution microscopy *(w-;Vps15-sgRNA#83822;Evi::GFP;hh-Gal4,UAS-Cas9uM/TM6B)* (n=4). Total Wg was stained (cyan), and endogenously tagged Evi/Wls::GFP (magenta) was used to analyse internalized Evi/Wls (Evi^In^). Regions shown at higher magnification are indicated by dashed boxes. **(f-f″)** In the wildtype half of the disc, Evi^In^ (magenta arrowheads) does not colocalize with total Wg (cyan arrowheads), although Evi/Wls is efficiently internalized in Wg-secreting cells. **(g-g″)** In Vps15-perturbed tissue, Evi^In^ (magenta arrowheads) is strongly reduced and mostly absent, while total Wg (cyan arrowheads) shows pronounced accumulation. The accumulated Wg signal does not overlap with any remaining Evi^In^ signal. **(h-h’’’)** Close-up of wild-type tissue showing total Wg (cyan) and Evi::GFP (magenta) staining. In the cytoplasm, areas of close proximity and colocalization between the two proteins are observed. At the apical membrane, colocalization is less apparent. Separate Wg signals (cyan arrowheads) are visible in the cytoplasm, as are distinct Evi/Wls signals (magenta arrowheads). **(e-h)** The predicted apical membrane region is marked with a horizontal dashed line. Confocal images were taken with the 63x oil objective and the YX view is presented as maximum intensity projection, while the ZX view are single slices. Imaging parameters for the super-resolution data can be found in Suppl. Table S5. All discs are oriented with anterior towards the left and perturbed areas are marked with vertical dashed lines. AFI: Average fluorescence intensities; mut: mutated.

Next, we asked whether Evi/Wls is still endocytosed from the apical membrane upon Vps15 perturbation. We therefore performed the same uptake assay described above, but instead of detecting internalized Wg, we monitored endocytosed Evi/Wls using an endogenously tagged Evi/Wls::GFP construct. To this end, we recombined Evi/Wls::GFP into our CRISPR/Cas9 driver fly line and crossed this line to the Vps15-sgRNA#83822 line. The uptake assay was performed as described by Witte et al. (2021), and in parallel total Wg was stained. In wildtype tissue, Evi/Wls was efficiently endocytosed from the apical membrane, as revealed by intracellular Evi/Wls::GFP (Evi^In^) signal (Fig. 5e; n=4). In contrast, strongly reduced Evi^In^ signal was detected in the Vps15 perturbed compartment. Higher-magnification of super-resolution imaging further showed that Evi^In^ does not colocalize with total Wg in wildtype tissue (Fig. 5f-f’’), suggesting that Wg and Evi/Wls are endocytosed separately in Wg-secreting cells. In the perturbed tissue, only weak and irregular Evi^In^ signals were observed (Fig. 5g-g″), which differed in morphology from those seen in wildtype cells.

Further, our super-resolution data provided a more detailed view of the localization of Evi/Wls and Wg during Wg secretion. In wildtype tissue, the Evi^In^ signal did not overlap with total Wg staining (Fig. 5f-f’’). Similarly, in the perturbed half, where some residual signal remained, no close proximity was observed. Additional analysis of wildtype tissue using super-resolution imaging with staining for total Wg and Evi::GFP (Fig. 5h-h’’’) revealed a mixed population of events: in some cases, Evi/Wls was in close proximity with Wg in the cytoplasm (white overlap of signals), while in other cases, Wg and Evi/Wls signals appeared separately without overlap. Notably, at the apical membrane, colocalization was rare and both proteins appear mainly as distinct signals. Together with the absence of colocalization following internalization (Evi^In^), these observations suggest that a substantial portion of the Wg-Evi/Wls complex dissociates at the apical membrane, prior to endocytosis.

### PI3K (III) and Evi/Wls post-endocytosis dynamics

In the previously shown Evi uptake assay, we observed a strong reduction of the Evi^In^ signal in the Vps15-perturbed tissue. However, it remains puzzling what exactly is happening to Evi/Wls in this tissue. The uptake assay could indicate that either the endocytosis of Evi/Wls is already reduced in the perturbed tissue or that the Evi-loaded endocytic vesicles undergo a different fate then Wg-loaded vesicles. It would be important to understand if endocytosis of Evi/Wls is impaired in Vps15 perturbed tissue or what the fate of the Evi-loaded endocytic vesicles is (Fig. 6b).

**Figure 6.**
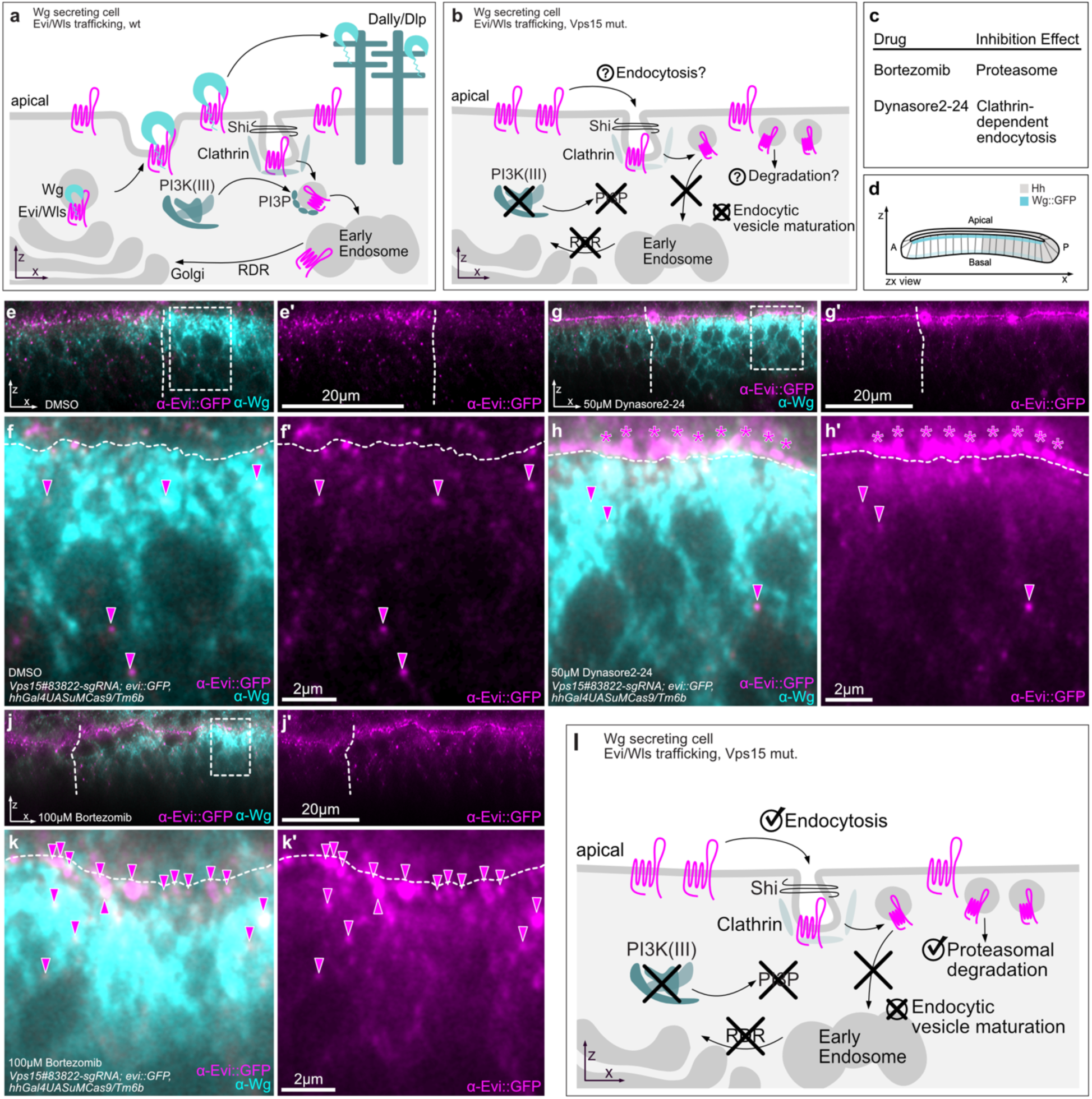
Proteasome-dependent degradation of Evi/Wls upon PI3K (III) perturbation in Wg-secreting cells. **(a)** Overview of Wg secretion in wing imaginal discs. Evi/Wls transports Wg to the apical membrane, where Wg is released for further trafficking. Evi/Wls is internalized from the apical membrane in a clathrin- and dynamin-dependent manner and trafficked to the endosomal pathway, followed by retromer-dependent recycling (RDR). **(b)** Working model of Evi/Wls dynamics upon Vps15 perturbation and loss of PI3P. Based on prior findings, it remained unclear whether PI3K (III) disruption affects Evi/Wls endocytosis (question mark). We further hypothesized that reduced Evi/Wls levels result from rapid proteasome-dependent degradation (second question mark) of Evi/Wls-loaded endocytic vesicles. **(c)** Overview of chemical inhibitors used in this study. **(d)** Schematic representation of the ZX view orientation shown in this figure. **(e-e’)** Control treatment with DMSO in SchM+ medium (n=5) in *w-;Vps15-sgRNA#83822;Evi::GFP;hh-Gal4,UAS-Cas9uM/TM6B* tissue. Evi^In^ (magenta) distribution in the wildtype compartment is comparable to previous observations. **(f-f’)** Higher magnification of the Vps15-perturbed compartment reveals only sparse Evi^In^ signal (magenta arrowheads), indicating residual endocytic activity. **(g-g’)** Treatment with the dynamin inhibitor Dynasore2-24 (50 µM) combined with the Evi/Wls uptake assay results in strong apical accumulation of Evi^In^ (magenta; n=10) in both wildtype and Vps15-perturbed compartments. **(h-h’)** Higher magnification of the perturbed region shows limited residual Evi^In^ signal (magenta arrowheads), suggesting incomplete inhibition of endocytosis. **(j-j’)** Treatment with the proteasome inhibitor Bortezomib (100 µM) in combination with the Evi/Wls uptake assay leads to accumulation of Evi^In^ (magenta; n=8). **(k-k’)** Higher magnification of the Vps15-perturbed compartment reveals pronounced apical Evi^In^ accumulation (magenta arrowheads). **(e-k)** Total Wg staining is shown in cyan. **(l)** Model summary: Upon Vps15 perturbation, Evi/Wls is still endocytosed but fails to enter the endosomal pathway. Instead, Evi/Wls-loaded endocytic vesicles are targeted for proteasome-dependent degradation. Impaired endosomal progression also disrupts retromer-dependent recycling, potentially contributing to the overall reduction of Evi/Wls levels in Wg-secreting cells. mut: mutated; Wt: Wildtype.

To address these questions, we employed pharmacological perturbations to dissect Evi/Wls trafficking. To understand if Evi/Wls reduction is due to an impaired endocytosis we tested an inhibitor which is targeting, according to the literature, the endocytosis of Evi/Wls in Wg secreting cells (Belenkaya et al., 2008; Franch-Marro et al., 2008; Gasnereau et al., 2011; Port et al., 2008). We applied the dynamin inhibitor Dynasore2-24 (Gagliardi et al., 2014; Kunduri and Acharya, 2024; Lalioti et al., 2025) to block endocytosis and test whether inhibition of internalization phenocopies the Vps15 defect. In a further experiment, we treated discs with the proteasome inhibitor Bortezomib (Velentzas et al., 2013) to assess whether endocytosed Evi/Wls might be degraded (Fig. 6c). We hypothesise at this moment, that due the role of PI3P in the endosomal maturation process, degradation via the lysosome which would depend on the endosomal pathway might not be likely (Piper and Luzio, 2007; Sandoval and Bakke, 1994). Both treatments were combined with the Evi/Wls uptake assay and analysed by super-resolution microscopy.

As a control, discs were treated with DMSO and subjected to the uptake assay (Fig. 6e–e′; n=3). As expected, Evi^In^ signal was strongly reduced in the Vps15-perturbed compartment, while endocytosis remained unaffected in the wildtype half (Fig. 6f-f’). Upon Dynasore2-24 treatment, a pronounced accumulation of Evi^In^ at the apical membrane was observed in both wildtype and Vps15-perturbed tissue (Fig. 6g–g′; n=10), indicating that Evi/Wls internalization is dynamin-dependent. Furthermore, this pronounced apical accumulation is in contrast to the reduction of Evi/Wls observed upon Vps15 perturbation and suggests that Evi/Wls endocytosis remains functional in Vps15-perturbed cells and that the phenotype is unlikely to arise from defective internalization.

Given that Evi/Wls endocytosis remains intact upon Vps15 perturbation, we next asked whether Evi/Wls-loaded endocytic vesicles are instead degraded. One possibility is proteasome-mediated degradation in the apical region of the disc proper. To test this, we treated wing discs with the proteasome inhibitor Bortezomib and performed the Evi/Wls uptake assay. Upon proteasomal inhibition, we observed a clear accumulation of Evi^In^ in the Vps15-perturbed tissue (Fig. 6j–j′; n=3). Higher magnification confirmed pronounced apical Evi^In^ accumulation under these conditions (Fig. 6k-k’). Together, these findings indicate that upon loss of PI3K (III) activity Evi/Wls-loaded endocytic vesicles are likely to be subject to proteasome-dependent degradation (summarized in Fig. 6l).

## METHODS

### Fly husbandry and fly lines

All fly lines were kept at 25°C with a 12/12-hour light/dark cycle. For the CRISPR-Cas9 screen we used the HD_CFD sgRNA library, which are available from the Vienna *Drosophila* Resource Centre (VDRC) (Port et al., 2020). For a full list of all screened lines, see Supplementary Table S1. The HD_CFD lines, containing the gRNAs, were crossed to a master stock line expressing Cas9 and two endogenously tagged read-out genes: *w-;Wg::GFP,Dll::dTom;hh-Gal4,UAS-Cas9uM/TM6b*. Other driver lines used in this study were *w-;Wg::GFP;hh-Gal4,UAS-Cas9uM/TM6b*, *w-;;Gal4-pdm2,UAS-Cas9uM/TM6b*, *w-;;hh-Gal4,UAS-Cas9uM/TM6b, w-;;act-Gal4,UAS-Cas9uM/TM6b* and *w-;;Evi-Wls::GFP,hh-Gal4,UAS-Cas9uM/TM6b*. The Wg::GFP fly line was generated by Port et al., 2014 and the Dll::dTom by Chaudhary et al., 2019. The Evi::GFP fly line was generated in our laboratory by B. Pavlovic and not published previously. Stocks obtained from the Bloomington *Drosophila* Stock Centre, BDSC (NIH P40OD018537) were used in this study. The following fly lines were obtained from BDSC: gRNA Vps15 (BDSC #83822), gRNA Atg6 (BDSC #80830), gRNA Vps34 (BDSC#92484), UAS Vps15 RNAi (BDSC #34029), UAS Vps15 RNAi (BDSC #57011), UAS Vps15 RNAi (BDSC #35209), UAS Atg6 RNAi (BDSC #102460), UAS Vps34 RNAi (BDSC #33384), UAS Vps34 RNAi (BDSC #36056), *nos-Cas9* (BDSC #54591) and UAS-FYVE-GFP (BDSC #42712).

### Fly genome editing and transgenesis

Endogenous fluorescent tagging of Evi/Wls locus was achieved using CRISPR/Cas9 homology-directed repair (Port et al., 2014). For this target site, a donor plasmid and a gRNA plasmid were generated. Donor plasmids contained the fluorescent tag flanked by 5′ and 3′ homology arms corresponding to the genomic regions adjacent to the insertion site. Evi/Wls donor plasmids were assembled in three steps: insertion of the homology arm, fluorescent protein coding sequence, and the opposite homology arm, using Type IIS restriction enzymes (BbsI or Esp3I; NEB) to avoid residual restriction sites. Homology arms were PCR-amplified from *Drosophila w1118* genomic DNA using the following primer sequences:

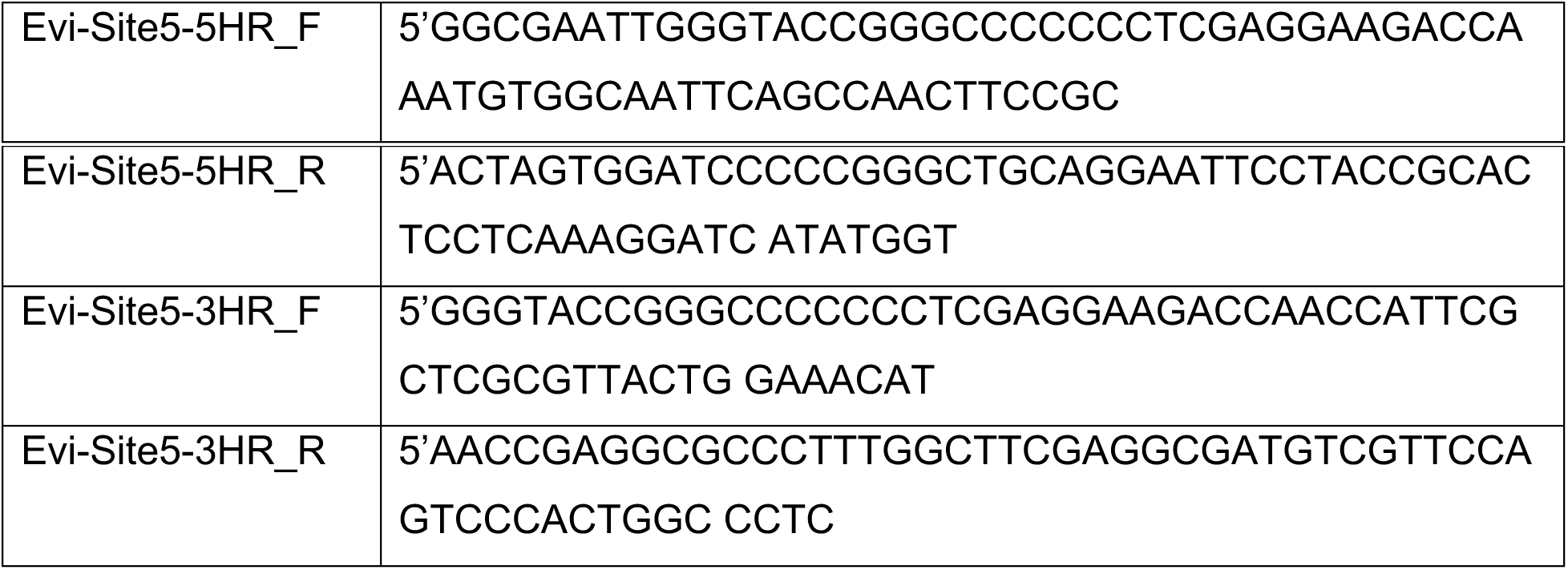

gRNA plasmids for Evi were constructed using the pCFD5 backbone (Port and Bullock, 2016), with sgRNAs selected adjacent to NGG PAM sequences (sgRNA1: 5’-TGCAGTGGGACTGGAACGACAATG and sgRNA2: 5’- AAACCATTGTCGTTCCAGTCCCAC). Oligonucleotide inserts were annealed and ligated into BbsI-digested pCFD5. Donor and gRNA plasmids were co-injected into posterior poles of *y,nos-Cas9,w-;;* embryos at 250 ng/µL. Embryos were collected, dechorionated in 10% bleach, aligned on glue-coated coverslips, covered with oil, and injected. Injected embryos were incubated at 18 °C in a humid chamber until hatching. Surviving larvae were transferred to standard food and reared at 25 °C. F1 progeny were screened by PCR using insert-specific primers and positive hits were crossed to balancers. The insertion was verified in stable stocks by PCR across homology arms and Sanger sequencing.

### Antibodies

For immunohistochemistry, the following primary antibodies were used: mouse anti-Wg (4D4, DSHB), 1:100 for immunofluorescence (IF) and 1:20 for extracellular staining; rabbit anti-Evi/Wls (Port et al., 2008), 1:800 for IF; rabbit anti-GFP (A11122, Invitrogen), 1:100-1:200 for IF and 1:20 for uptake assay; mouse anti-Dlg1 (4F3, DSHB), 1:50 for IF; mouse anti-Dlp (13G8, DSHB) 1:200 for IF. As secondary antibodies we used the following antibodies at 1:800 dilution: chicken anti-mouse Alexa Fluor 594 (A21201, Invitrogen), goat anti-rabbit Alexa Fluor 594 (A11012, Invitrogen), goat anti-mouse Alexa Fluor 647 (A21236, Invitrogen), goat anti-rabbit Alexa Fluor 647 (A32733, Invitrogen), goat anti-guinea pig Alexa Fluor 594 (A11076, Invitrogen). For STED super resolution microscopy, we used the secondary antibodies goat anti-rabbit STAR 635P (ST635P-1002, Abberior) and goat anti-mouse STAR 635P (ST635P-1001, Abberior) in 1:200 dilution. All imaginal discs were additionally stained with Hoechst33342 (H1399, Invitrogen) in 1:2000 dilution.

### Sanger Sequencing

To verify the efficiency of the CRISPR-Cas9 sgRNA lines used in this study, we crossed the sgRNA lines sgRNA-Vps15#009 (sgRNA: 5’-ACAAGGCAGCCTACATAATG and XXX) and sgRNA-Vps15#83822 (sgRNA: 5’-CAAGGCAGCCTACATAATGC) to *w-;;act- Gal4,UAS-Cas9* flies to achieve ubiquitous Cas9 expression throughout larval tissues. For each cross, ten third instar larvae were collected, and genomic DNA was extracted of each larvae using squishing buffer (10 mM Tris-HCl pH 8.2, 1 mM EDTA, 25 mM NaCl) supplemented with freshly added Proteinase K (200 µg/mL). For Sanger sequencing, the Vps15 locus was PCR-amplified for each single larvae using the primers forward 5′-GCCACCAGCTGTTTGGCTGC and reverse 5′-CTCCACCAGATCTCGCTCGC with 2X DreamTaq Green Master Mix (Thermo Scientific, K1081). PCR products were purified using the NucleoSpin Gel and PCR Clean-up Kit (Macherey-Nagel) and separately sequenced using the forward primer. Sequencing traces were analysed for each larvae using the EditCo ICE (Inference of CRISPR Edits) tool (v3.0; https://ice.editco.bio). A detailed summary of editing efficiencies is provided in Supplementary Table S4.

### Immunohistochemistry

Imaginal wing discs of third instar larvae (wandering stage) were dissected according to standard procedures. Wing discs were fixed with 4% formaldehyde (FA) in 1X PBS for 30’ and washed several times with 1X PBS-Triton X-100 0.1% (PBST). For the CRISPR screen using endogenously labelled Wg::GFP and Dll::dTom, discs were mounted in anti-fade mounting medium VectaShield plus DAPI (VectorLabs) after several times washed with 1X PBST. For immunofluorescence, discs were blocked using 1% normal goat serum in PBS for 1h and incubated overnight at 4°C with the primary antibody in 1X PBS. On the following day, discs were washed several times with 1X PBST and either incubated for 2h with the secondary antibodies in 1X PBST or overnight at 4°C with the secondary antibody in 1X PBS. After incubation with the secondary antibody, discs were washed several times with 1X PBST and mounted in VectaShield (VectorLabs) on Poly-L-Lysin (P1524, Sigma-Aldrich) coated cover slips.

For extracellular Wingless staining, larvae were kept for 15’ prior to dissection in ice cold Schneider’s insect medium (S0146, Sigma) with 4% FBS (SchM+) and subsequently dissected on ice in SchM+. Staining was performed with incubation on ice for 90’, with the primary antibody (1:20 with anti-Wg (4D4, DSHB)) in SchM+. Discs were washed with SchM+ and fixed with 4% FA in SchM+ on ice for 40’, followed by another 10’ fixation at room temperature. After these steps, discs were treated with the above-described immunohistochemistry procedures. Here, we incubated the discs additionally with rabbit anti-GFP primary antibody to stain the total amount of Wg additionally to the extracellular Wg (detected by mouse anti-Wg). For confocal microscopy we used the secondary antibodies goat anti-mouse Alexa Fluor 647 (A21236, Invitrogen) for extracellular Wg staining and goat anti-rabbit Alexa Fluor 594 (A11012, Invitrogen) for total Wg::GFP detection at a 1:200 dilution.

The Wg uptake assay was performed according to procedures described in (Witte et al., 2021). Wing discs were dissected at room temperature in SchM+ and incubated in SchM+ with the primary antibody for 90’. We used as primary antibodies either 1:20 of mouse anti-Wg, (4D4) or 1:20 of rabbit anti-GFP (A11122, Invitrogen) for the Evi/Wls Uptake assay detecting Evi::GFP. The reaction was stopped using an acid wash with 0.1mM Glycine with HCl (pH 3.5) for 5’, followed by stringent washes with SchM+ (5 times, 3-5’). After washing, discs were fixed for 30’ at room temperature in 4% FA in SchM+. After fixation, discs were washed with 1X PBS and treated with a normal immunohistochemistry procedure. Additionally, to the incubation with primary antibody during the Uptake assay, we incubated the discs after fixation with another primary antibody. For Wg Uptake (mouse anti-Wg) we used additional rabbit anti-GFP antibody for total Wg staining. For Evi/Wls Uptake (rabbit anti-GFP) we used additionally mouse anti-Wg (4D4) also for detection of total Wg. For confocal microscopy of Wg Uptake, we used the secondary antibodies goat anti-mouse Alexa Fluor 647 (A21236, Invitrogen) and goat anti-rabbit Alexa Fluor 594 (A11012, Invitrogen). For Evi/Wls Uptake we used chicken anti-mouse Alexa Fluor 594 (A21201, Invitrogen) and goat anti-rabbit Alexa Fluor 647 (A32733, Invitrogen). For STED microscopy of Wg Uptake, we used goat anti-mouse STAR 635P (ST635P-1001, Abberior) for detecting endocytosed Wg and goat anti-rabbit Alexa Fluor 594 (A11012, Invitrogen) for total Wg staining. For the Evi/Wls Uptake we used goat anti-rabbit STAR 635P (ST635P-1002, Abberior) for endocytosed Evi/Wls detection and chicken anti-mouse Alexa Fluor 594 (A21201, Invitrogen) for total Wg staining. All secondary antibodies were used at a 1:200 dilution. All discs for confocal microscopy were mounted in the anti-fade mounting medium VectaShield (VectorLabs) and placed on a Poly-L-Lysin (P1524, Sigma-Aldrich) covered cover slip. For mounting discs for STED microscopy please refer to separate subsection below.

### Ex vivo inhibitor treatment

For the treatment with cytotoxic drugs, third instar larvae were collected and dissected in SchM+ similar to the Uptake assay. For proteasomal inhibition, imaginal wing disc were incubated with 100μM Bortezomib in DMSO (PS-341; S1013, Selleckchem; CAS: 179324-69-7) in SchM+ for 90’. After incubation discs were washed briefly with SchM+ and fixed in 4% FA in SchM+ for 30’ at room temperature. For inhibition of endocytosis, wing discs were incubated with 50μM Dynasore2-24 Dyngo-4a (also known as Dyngo-4a; HY-13863; MCE; Cas: 1256493-34-1) and treated similar as described above. As negative control, discs were incubated with SchM+ containing DMSO (1:1000; Sigma). After fixation, discs were fluorescently stained using mouse anti-Wg (4D4, DSHB) and rabbit anti-GFP (A11122, Invitrogen) primary antibodies. For STED microscopy, the following secondary antibodies were used: chicken anti-mouse Alexa Fluor 594 (A21201, Invitrogen) and goat anti-rabbit STAR 635P (ST635P-1002, Abberior). The same procedure was used when this treatment was combined with Evi/Wls uptake assay. In this case, discs were incubated with the primary anti-GFP (A11122, Invitrogen) antibody simultaneously with the drug treatments described above. Incubation was carried out for 90’ in SchM+ medium. Notably, the uptake assay was terminated by an acidic wash prior to fixation. All subsequent steps were identical to those described for the *ex vivo* inhibitor treatment.

### Fluorescent in situ hybridisation (FISH)

For *in situ* hybridisation, L3 wing imaginal discs were dissected in 1X PBS and fixed for 2x 20 min in 4% FA in PBS or PBST respectively. Tissues were hybridised at 65°C overnight with DIG-labelled *wg* antisense probe in hybridisation buffer HybA (50% formamide, 5x SSC pH 7.0, 10 mg/mL salmon sperm, 1 mg/mL tRNA, 100 μg/mL heparin and 0.2% Triton X-100, adjust pH 6.5 with 1M HCl). *In situ* antisense probes were generated using a two-step PCR based method with the *wg* specific primers forward 5’-CTCCCGGGAATTCGTCGATA, reverse 5’-TTTTGGTCCGACACAGCTTG and a universal 3’ T7 primer 5’agggatcctaatacgactcactatagggcccggggc3’. Probes were transcribed with RNA T7 polymerase (EP0113, Thermo Scientific) and labelled with the DIG RNA labelling mix (#11277073910, Roche). After hybridisation, tissues were washed with 1X PBST and blocked with 1X blocking reagent (#11096176001, Roche) in 1X Maleic acid buffer (100mM Maleic acid, 150mM NaCl, pH7.5). Samples were incubated overnight with the HRP-conjugated anti-DIG antibody (1:2000, Jackson, AB_2339011) in 1X blocking reagent (Roche). For fluorescent signal detection, samples were incubated with 1:50 1X TSA amplification diluent (FP1135, AKOYA bioscience) and CF555 conjugated tyramide (#96021, Biotium Inc.) for reaction with HRP-conjugated antibody. Wing discs were mounted on Poly-L-Lysin (P1524, Sigma-Aldrich) coated coverslips and embedded in VectaShield anti-fade mounting medium (VectorLabs).

### Light Microscopy and Super resolution microscopy

Wing discs were imaged using a Leica TCS SP8 confocal microscope (Leica Microsystems) equipped with a 40x/1.30 HC PL APO CS2 (oil) and 63x/1.40 (oil) objectives. Brightfield images of adult wings were acquired using a Zeiss Cell Observer.Z1 microscope with a 20x/0.8 Plan Apo DICII objective. These images were captured using an AxioCam MRm monochrome CCD camera (Carl Zeiss Microscopy GmbH). 3D-STED super resolution images were acquired with near isotropic resolution (Klar et al., 2000) as ZX-scans with a Leica Stellaris 8 STED Falcon microscope (Leica Microsystems). Imaginal discs were prepared and mounted in 1X PBS on a Poly-L-Lysin (P1524, Sigma-Aldrich) covered μ-Dish (35mm) with a high 1.5H glass coverslip bottom (#81158, Ibidi) and covered with 1.5H round (012mm) coverslips (Marienfeld). Imaging was performed with a HC PL APO 86x/1.20 W motCORR STED white objective (Leica Microsystems) at a pixel size of about 0.04μm. For STED microscopy we used exclusively the following secondary antibody combinations: (i) chicken anti-mouse Alexa Fluor 594 (A21201, Invitrogen) and goat anti-rabbit STAR 635P (ST635P-1002, Abberior) or (ii) goat anti-rabbit Alexa Fluor 594 (A11012, Invitrogen) and goat anti-mouse STAR 635P (ST635P-1001, Abberior). All detailed imaging parameters of presented STED images can be found in Supplementary Table S5.

## Data processing

Image data was processed and analysed using Fiji ImageJ suit Version 2.1.0/1.53c (Schindelin et al., 2012). Further analysis and data visualisation was performed using RStudio Version 4.5.0 (R Core Team (2023), R: A Language and Environment for Statistical Computing, R Foundation for Statistical Computing, Vienna, Austria; RStudio Team (2020), RStudio: Integrated Development for R, RStudio, PBC, Boston, MA). In some cases, fluorescence intensity measurements were obtained from images acquired at different resolutions, resulting in varying pixel densities across the imaged regions. To enable direct comparison, spatial distances were normalized by scaling each profile to a maximum value of 1, achieved by dividing each distance value by the maximum distance per image. To account for variability in staining intensity between samples, pixel intensity values were also normalized. Assuming no biological differences at the edges of the imaged regions, the leftmost 10% of pixels were used as a baseline. The median intensity of these baseline pixels was calculated, and all intensity values were normalized by dividing by this median. Trend lines were fitted using the R function geom_smooth with the method set to “loess”. Statistical differences between perturbed and unperturbed regions were assessed using the Wilcoxon rank-sum test (wilcox.test) in R. Figures with images and schematic drawings were composed by Affinity Designer Version 1.10.8 (Serif).

## DISCUSSION

In this study, we identify class III PI3K activity as a key regulator of Wg secretion dynamics at the DV boundary of *Drosophila* wing imaginal discs. Using a CRISPR-Cas9 screen, we reveal that loss of the PI3K (III) subunit Vps15 causes a pronounced apical accumulation of Wg by selectively disrupting an endocytosis-dependent “indirect” Wg pool, while the direct, glypican-mediated Wg transport remains largely unaffected. This phenotype reflects a block in endocytic vesicle maturation. In contrast, the Wg cargo receptor Evi/Wls exhibits distinct trafficking behaviour: its levels are reduced upon Vps15 perturbation, likely due to proteasome-dependent degradation of Evi/Wls-loaded endocytic vesicles. Together, our findings define a specific role for PI3K (III) in post-endocytic Wg trafficking and uncover a previously unrecognized divergence between Wg and Evi/Wls dynamics at the apical membrane.

*In vivo* CRISPR screening combined with endogenously tagged readout genes proved to be a powerful approach for identifying regulators of Wg secretion in *Drosophila*. Our screen robustly recovered known Wnt pathway components, including *myopic (mop), microtubule star (mts), Pp1α-96A, myotubularin (Mtm),* and *Casein kinase Iα (Ck1α),* demonstrating strong concordance with previous RNAi-based studies (Swarup et al., 2015). In addition, we identified a small number of previously unrecognized candidates. Notably, Vps15 had been included in earlier RNAi screens but was not identified to play a role in Wg secretion (Chaudhary and Boutros, 2019; Gross et al., 2012; Port et al., 2011; Swarup et al., 2015), likely due to the weak or absent phenotype upon partial knockdown. In contrast, CRISPR-mediated mutagenesis revealed a robust Wg phenotype, highlighting the value of near-complete gene disruption for uncovering key regulators.

Loss of Vps15 function leads to a pronounced accumulation of Wg at the apical region of Wg-secreting cells along the DV boundary. This phenotype is fully phenocopied by perturbation of Atg6 or Vps34, the other core subunits of the class III PI3K complex, demonstrating that the observed Wg accumulation is a specific consequence of impaired PI3K (III) activity and requires the integrity of the entire complex. Although previous studies reported Wg accumulation following Atg6 knockdown using RNAi (Abe et al., 2009; Lőrincz et al., 2014), the underlying cellular mechanisms remained unclear. Furthermore, while a role for Vps34 in Wg receiving cells has been proposed, this model does not readily explain the phenotype observed in Wg-secreting cells (Hemalatha et al., 2016). Together, these findings indicate that, despite earlier insights into PI3K (III) function in *Drosophila*, its precise role in regulating Wg secretion and trafficking within Wg-secreting cells had not been fully resolved.

Our analysis of the long range Wg gradient (Wg^Ex^), combined with the Wg uptake assay and colocalization with Dlp, indicates that the direct Wg signalling pool and its transfer onto glypicans are not affected by loss of PI3K (III) function. This contrasts with phenotypes observed upon loss of Evi/Wls or Shibire (Dynamin) in Wg-secreting cells of the wing imaginal disc, which impair Wg secretion and gradient formation (Bänziger et al., 2006; Bartscherer et al., 2006; Strigini and Cohen, 2000). As shown by previous studies, PI3K (III) phosphorylates PI to generate PI3P, a lipid required for maturation of endocytic vesicles (He et al., 2017; Posor et al., 2022). Consistent with this role, we show that loss of PI3K (III) leads to the apical accumulation of Wg-loaded endocytic vesicles in the disc proper. Future application of advanced lipid-labelling techniques, such as lipid sensors or metabolic labelling (Hale et al., 2020; He et al., 2017) in *Drosophila* could provide direct evidence for the depletion of PI3P in these vesicles.

Under physiological conditions, “indirectly” trafficked Wg is routed through the endosomal system and can be further transported via multivesicular bodies onto exosomes, directed toward basal secretion, or conveyed through cytoneme-mediated mechanisms (Brunt and Scholpp, 2018; Gross, 2021; Gross et al., 2012; Moti et al., 2019; Stanganello and Scholpp, 2016). The specific trafficking impacts of Vps15 knockout on this “indirect” Wg pool remain to be elucidated. Future studies could examine whether exosome-associated or filopodia-mediated Wg transport is altered in this tissue (Gross et al., 2012; Panáková et al., 2005). Moreover, Vps15 perturbation, together with our toolkit of endogenously tagged proteins and HD_CFD driver lines, may provide a valuable framework to further dissect alternative Wg trafficking routes in Wg-secreting cells in *Drosophila*.

In contrast to Wg, the cargo receptor Evi/Wls exhibits a distinct response to Vps15 perturbation. Loss of Vps15 causes a significant reduction in Evi/Wls protein levels. A similar phenotype has been reported upon loss of retromer function, where Evi/Wls levels decrease specifically in Wg-secreting cells (Belenkaya et al., 2008; Franch-Marro et al., 2008; Port et al., 2008). In that context, impaired retromer-dependent recycling and potential lysosomal degradation were suggested to underlie the observed reduction, highlighting the importance of proper recycling for maintaining the overall Evi/Wls abundance. To investigate Evi/Wls dynamics under Vps15 perturbation, we performed an Evi/Wls uptake assay. This assay alone did not conclusively reveal whether endocytosis of Evi/Wls is affected, or if, as with Wg, only the maturation of Evi/Wls-loaded endocytic vesicles is impaired. By combining the uptake assay with chemical inhibition of endocytosis and proteasomal degradation, we demonstrated that Evi/Wls is efficiently endocytosed but Evi-loaded endocytic vesicles are likely to be subsequently degraded, potentially via a proteasome-dependent mechanism. While a phenomenon like this where endosomal vesicles are targeted by proteasomal degradation, has been subject of discussion in other model organisms (Strous and Govers, 1999), it remains open if this mechanism is an emergency escape route of the cell to get rid of the excess of unbound Evi/Wls in the context of Vps15 perturbation or if it also occurs under wildtype conditions. Since proteasomal degradation of Evi/Wls has been described at the endoplasmic reticulum (Glaeser et al., 2018; Wolf et al., 2021), it is possible that similar targeting signals contribute to its degradation in the Vps15-perturbed state. Loss of Evi/Wls from the recycling pathway alone appears sufficient to reduce overall protein abundance (Belenkaya et al., 2008; Franch-Marro et al., 2008; Port et al., 2008), which could explain the phenotype observed upon Vps15 perturbation. Still, further studies are needed to clarify the degradation of Evi/Wls upon Vps15 perturbation.

This finding suggests that Wg- and Evi/Wls-loaded vesicles respond differently to Vps15 perturbation and that the effects of PI3K (III) loss act separately on these two distinct vesicle populations. Using 3D-STED super-resolution microscopy, we resolved the spatial relationship between Wg and Evi/Wls in wildtype and Vps15-perturbed tissue. While Wg- and Evi/Wls-positive particles are frequently in close proximity in the cytoplasm, they are not consistently colocalized at the apical membrane. Together with our uptake and chemical inhibition experiments, these observations suggest that the Wg-Evi/Wls complex begins to dissociate earlier than previously proposed (Coombs et al., 2010; Sharma and Chaudhary, 2024). Our data indicate that a substantial portion of the complex separates at the apical membrane, where Wg follows distinct “direct” or “indirect” secretion routes, and Evi/Wls is shuttled toward retromer-dependent recycling. This is consistent with recent findings showing that the lipid composition of the apical membrane itself is critical for Wg endocytosis and subsequent trafficking through the endosomal pathway (Alvarez-Rodrigo et al., 2026). We do not exclude additional dissociation events in late endosomes, but our results argue against this being the sole site of separation. If separation occurs at the apical membrane, important questions remain, including the molecular factors driving complex dissociation, the mechanisms guiding Wg onto glypicans, the determinants of the “direct” versus “indirect” Wg pool, and how Wg integrates into the membrane of endocytic vesicles.

In this study, we applied advanced imaging approaches to dissect Wg secretion dynamics in Wg-secreting cells at the DV boundary of the wing imaginal disc. 3D-STED super-resolution microscopy provided high-resolution insights into protein dynamics with cell-type specificity, revealing details that are not accessible with the resolution of conventional, diffraction-limited imaging. However, the narrow, columnar architecture of wing disc epithelial cells still limits precise subcellular localisation, making it challenging to assign proteins to specific organelles. Combining super-resolution microscopy for example with expansion microscopy (Zhuang and Shi, 2023) would represent a powerful strategy to overcome these limitations, offering the potential to visualise Wg and Evi/Wls trafficking with unprecedented spatial detail and to uncover new aspects of secretion and signalling mechanisms *in vivo*.

## CONCLUSIONS

In summary, our study uncovers distinct trafficking routes for Wg and its cargo receptor Evi/Wls in Wg-secreting cells. We show that PI3K (III) activity is essential for maturation of Wg- and Evi/Wls-loaded endocytic vesicles and their progression through the endosomal pathway. We demonstrated that upon PI3K (III) perturbation, the fate of these endocytic vesicles is differing depending on their cargo. While Wg-loaded vesicles are accumulating, Evi/Wls-loaded vesicles undergo proteasome-dependent degradation. Using super-resolution microscopy, we further demonstrate that the Wg-Evi/Wls complex is already separating at the apical membrane prior to endocytosis, creating two distinct endocytic vesicle populations. Beyond mechanistic insight, Vps15 perturbation provides a valuable experimental tool to dissect specific steps in Wg trafficking *in vivo*. These findings and emphasize the utility of super-resolution approaches to resolve trafficking events that are not accessible with conventional imaging. These findings highlight the complexity of Wg secretion and provide a clearer picture of the spatial dynamics of Wg secretion in Wg-secreting cells.

## LIST OF ABBREVIATIONS

A: Anterior
Act: Actin
AFI: Average Fluorescence intensity
Cas9: Caspase 9
CRISPR: Clustered Regularly Interspaced Short Palindromic Repeats
D: Dorsal
Dll: Distalless
Dlp: Dally-like protein
DMSO: Dimethyl sulfoxide
dTom: dTomato
DV: Dorso-ventral (boundary)
ECM: Extracellular matrix
ER: Endoplasmic Reticulum
Evi/Wls: Evenness-interrupted/Wntless
Evi^In^: Internalized Evi/Wls
GFP: Green Fluorescent protein
HD_CFD: Heidelberg CRISPR Fly Design (library)
hh: Hedgehog
Mut: Mutated
P: Posterior
Pdm2: POU domain protein 2
PI: Phosphoinositol
PI3K (III): Phosphoinositol 3 Kinase class III
PI3P: Phosphoinositol-3-phosphate
RNAi: RNA interference
ROI: Region of interest
sgRNA: small guide RNA
Shi: Shibire
UAS: Upstream activation sequence
V: Ventral
Wg: Wingless
Wg^Ex^: Extracellular Wg
Wg^In^: Internalized Wg
Wt: Wildtype

## DECLARATIONS

### Availability of data and materials

All relevant data and details of resources can be found within the article and its supplementary information. Original raw microscopy data is available upon request.

## Author’s contribution

MH: Designed Study, conducted experiments, acquired and analysed data, writing original draft, writing – review and editing; BP: conducted experiments, writing – review and editing; MM: conducted fly experiments; CS: conducted experiments and maintained fly stocks; JG: bioinformatics, writing – review and editing; FP: project planning, writing – review and editing; MB: funding acquisition, conceptualization, writing – review and editing.

## Acknowledgements

Research in the group of M.B. is supported by the Deutsche Forschungsgemeinschaft Collaborative Research Center CRC/SFB1324 Projects A01 (project number 331351713) on Mechanisms and Functions of Wnt signalling. The authors thank the Wnt CRC Consortium for the numerous stimulating discussions and exchanges. We would like to thank the DKFZ light microscopy core facility (LMCF) for their expertise and support.

