## Supplemental Information for "Class III PI3K is essential for Wingless secretion and Evi/Wls recycling in *Drosophila*"

### SUPPLEMENTARY INFORMATION

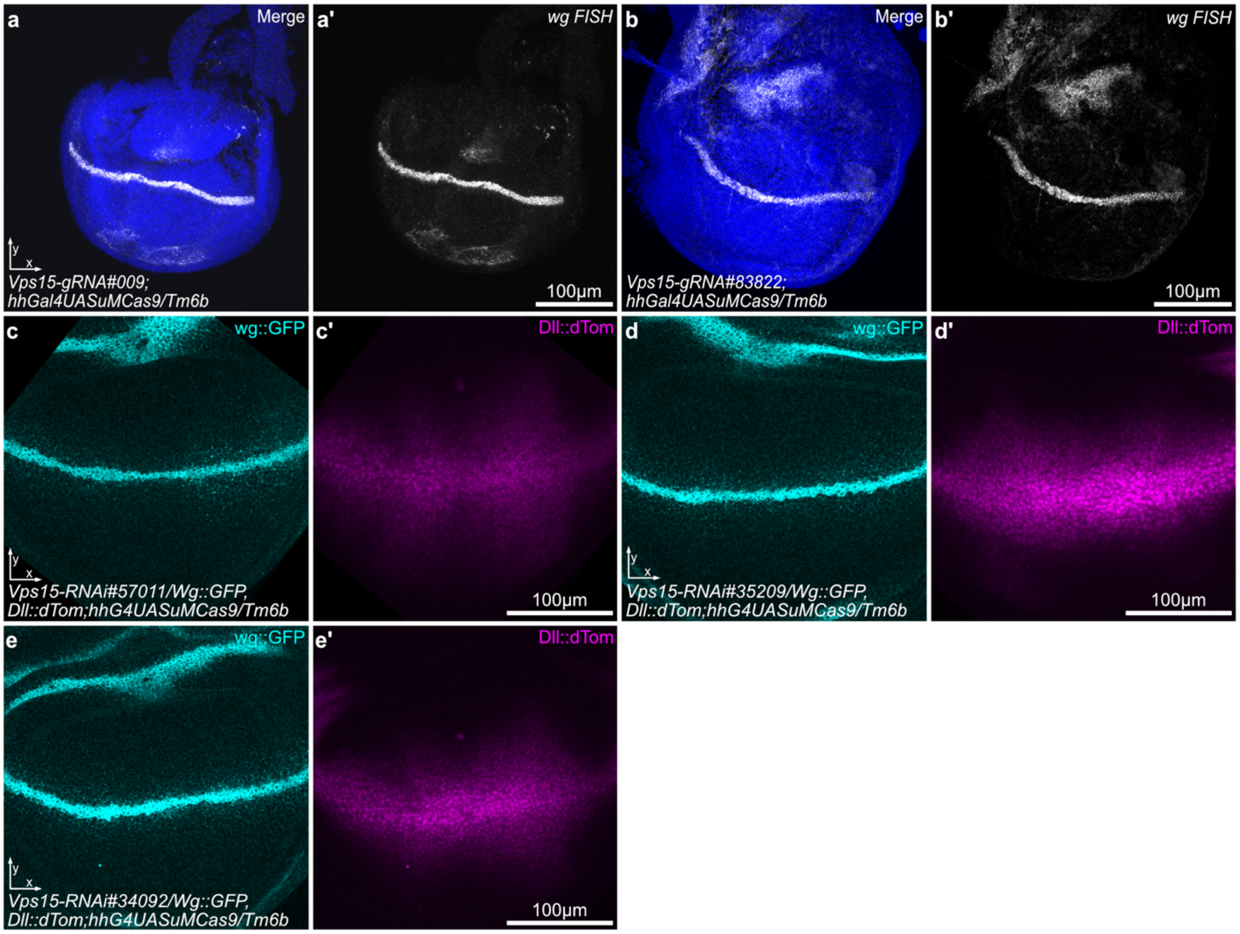


**Supplementary Figure S1 | Characterisation of Vps15 mutagenesis in the wing imaginal disc.** **(a,a')** Fluorescent in situ hybridisation of *Vps15-sgRNA#009* with the driver line *w-;;hhGal4UasCas9uM/Tm6b*. No increase in *wg* gene expression can be observed at the DV boundary. *wg* FISH signal in grey. **(b,b')** FISH of *Vps15-sgRNA#83822* with the driver line *w-;;hhGal4UasCas9uM/Tm6b*. No increase in *wg* gene expression can be observed at the DV boundary. *wg* FISH signal in white and Hoechst33342 in blue. **(c-e)** Knockdown of Vps15 in the wing disc with three different RNAi lines did not show any effect on Wg (cyan) or Dll (magenta) at the DV boundary. **(c,c')** BDSC Vps15 RNAi #57011 line, **(d,d')** Vps15 RNAi #35209 and **(e,e')** Vps15 RNAi #34092. **(a,b)** Images taken with 40x objective and **(c-e)** images taken with 63x objective. All confocal images are presented as maximum intensity projections. All discs are additionally stained with Hoechst33342 (blue).

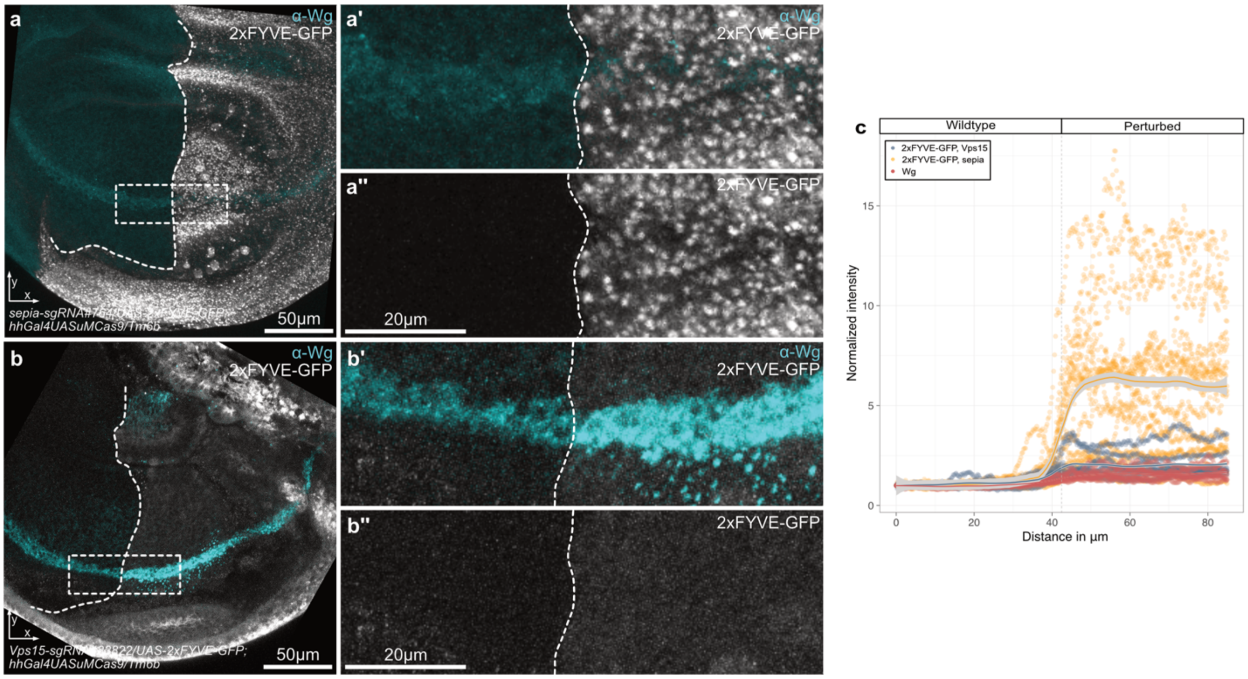


**Supplementary Figure S2 | Loss of 2xFYVE-GFP signal confirms functional disruption of PI3K(III).** **(a-a'')** Control cross using sepia-sgRNA (n=15) and the hhGal4, UAS-Cas9 driver combined with UAS-2xFYVE-GFP to label FYVE-domain containing vesicles. A strong GFP signal (grey) is observed within the hh expression domain. Total Wg staining is shown in cyan. **(a',a'')** Higher magnification of the dorsoventral (DV) boundary. **(b-b'')** Perturbation of Vps15-sgRNA #83822 (n=8) using the same system drastically reduces the GFP signal (grey) in the hh domain of the wing disc. Increased accumulation of Wg (cyan) is observed in the posterior compartment upon Vps15 perturbation. **(c)** Quantification of average UAS-2xFYVE-GFP fluorescence intensity. For reference, total Wg intensity (red) in Vps15-perturbed discs is also shown. GFP signal is indicated in blue for Vps15-perturbed cells and in yellow for sepia controls. The quantification mirrors the visual observation of a strong reduction in GFP signal upon Vps15 perturbation, confirming functional loss of PI3K(III) activity. All confocal images were taken with the 63x oil objective. Presented are maximum intensity projections for the YX view.

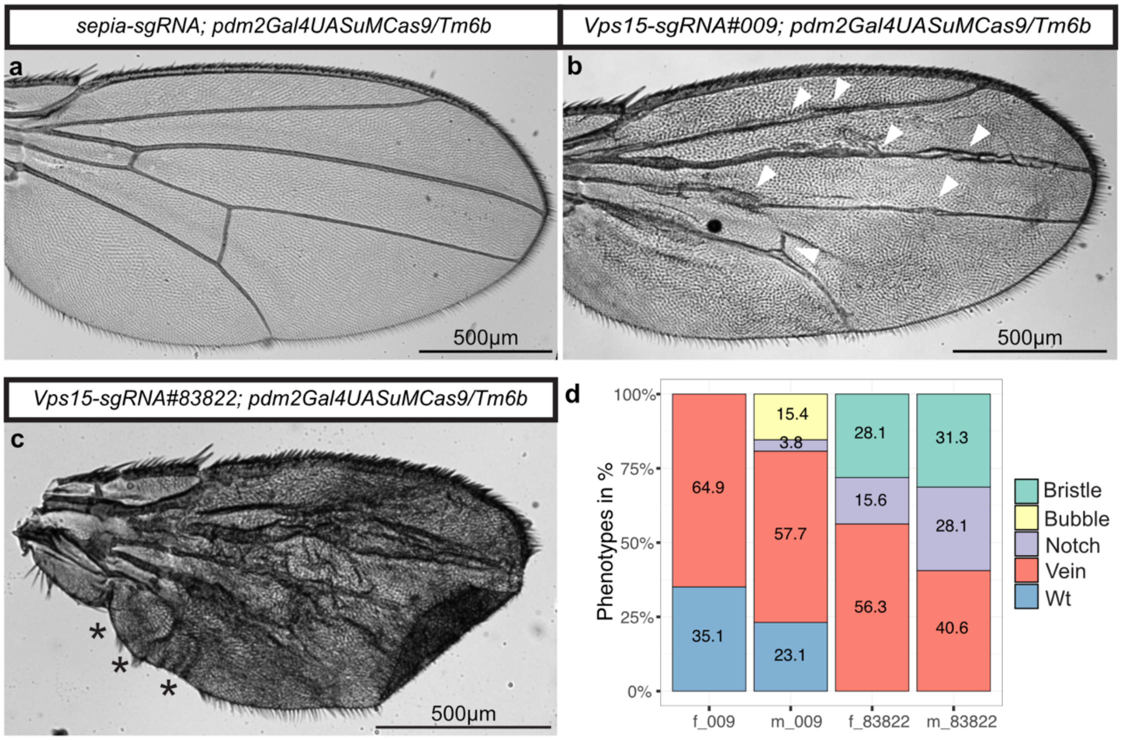


**Supplementary Figure S3 | Effects of Vps15 perturbation on adult *Drosophila* wings. (a-c)** Adult wings of crosses with the *w-; ;pdm2-Gal4,UASCas9uM/Tm6B* with **(a)** *sepia-sgRNA* as a negative control indicating the wildtype wing and **(b)** with *Vps15-sgRNA#009* or **(c)** *Vps15-sgRNA#838322* showing the perturbed wing tissue. **(b)** *Vps15-sgRNA#009* perturbed wings show a vein phenotype (white arrowheads). Further phenotypes observed include bubbles or notches, both also combined with vein phenotypes (Pictures not shown). **(c)** Wings with *Vps15-sgRNA#83822* perturbations show a more drastic phenotype than wings in (b). All wing tissue is highly disrupted while notches (black asterisks) and bristle phenotypes are observed regularly. Scale bar = 500um. **(d)** Distribution of wing phenotypes in *Vps-sgRNA#009* female ($n_{female}$=36), male ($n_{male}$=24) and *Vps15-sgRNA#83822* female ($n_{female}$=32), male ($n_{male}$=32) mutated flies. In *Vps15-sgRNA#009* flies wildtype wings were observed in female and male samples, while most wings show a vein phenotype. In male wings, notches and bubbles also occurred. No wildtype wings occurred in *Vps15-sgRNA#83822* female and male samples. All wings are disrupted and show a vein phenotype. Additionally notches and loss of bristles are observed in both sexes. Numbers within the coloured bars indicate percentages of each corresponding phenotype. f = female; m = male; Wt: wildtype.

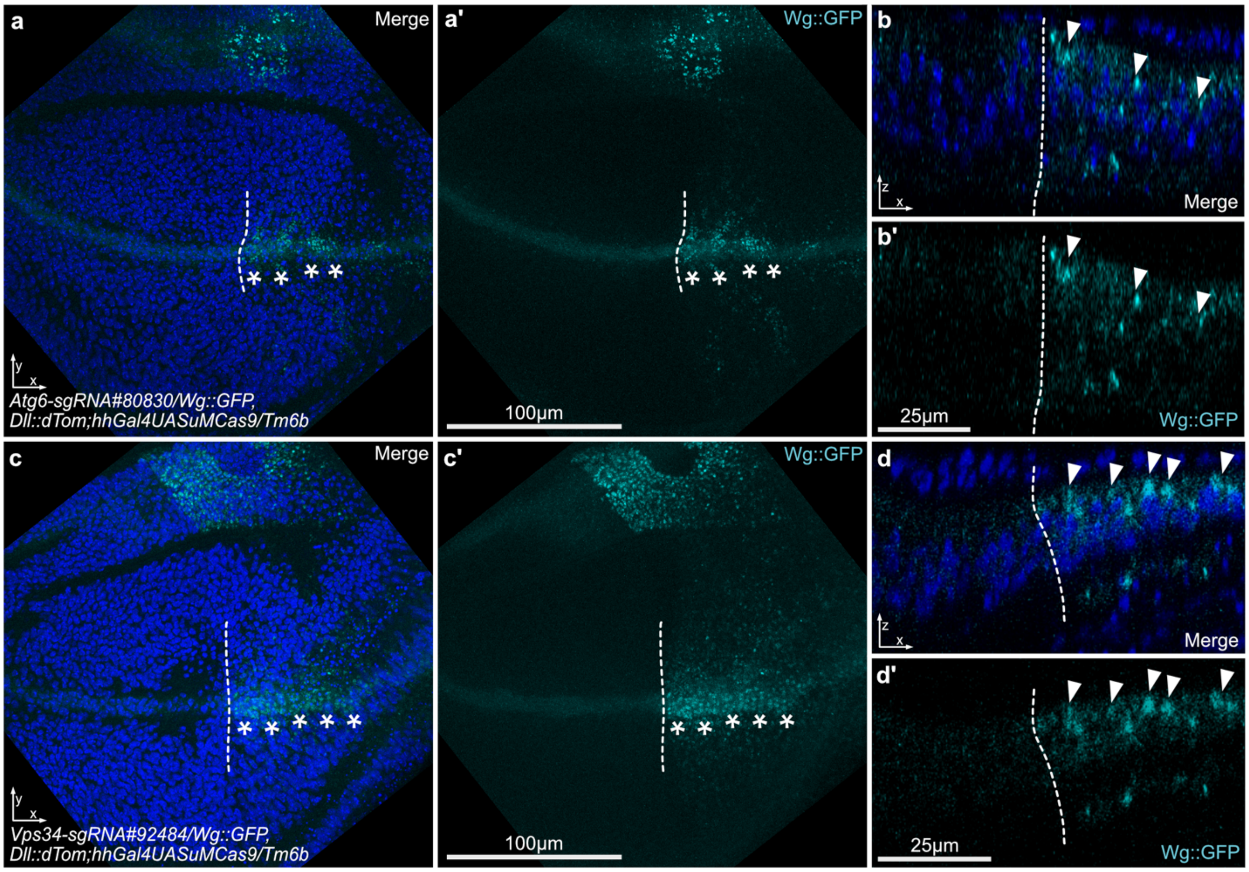


**Supplementary Figure S4 | Phenotypic characterisation of Atg6 and Vps34 mutagenesis in the wing imaginal disc. (a,a')** Wg protein abundance at the DV boundary in the wing discs of *w-; Atg6-sgRNA#80830, Wg::GFP,Dll:dTom;hh-Gal4,UASCas9uM/Tm6B*. Atg6 mutagenesis shows accumulation of Wg protein (cyan) at the DV boundary (asterisks). **(b,b')** Wg accumulation (cyan) is also detected at the apical membrane in the ZX view (arrowheads). **(c,c')** Wing imaginal disc of the cross of *w-; Vps34-sgRNA#92484, Wg::GFP,Dll:dTom;hh-Gal4,UASCas9uM/Tm6B* shows accumulation of Wg::GFP at the DV boundary (asterisks). The loss of Vps34 phenocopies Atg6 and Vps15 phenotypes. Wg protein (cyan) accumulation at the DV boundary can be detected and **(d,d')** this Wg (cyan) accumulation is also localised at the apical membrane. Nuclei were stained with Hoechst33342 (blue). All discs are orientated with the wildtype tissue towards the left (anterior) and perturbed tissue towards the right (posterior). If possible, borders between both regions are indicated with white dashed lines. All confocal images were taken with the 63x oil objective. Presented are maximum intensity projections for the YX view and single slices for the ZX view.

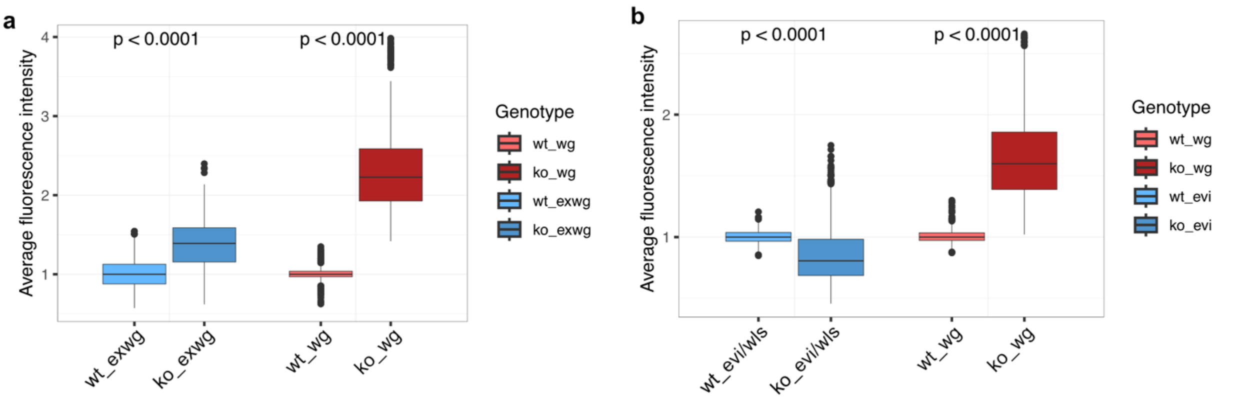


**Supplementary Figure S5 | Statistical tests of fluorescence measurements for Wg$^{Ex}$ and Evi/Wls.** Shown are boxplots corresponding to data shown in Fig. 3b and Fig. 5c. **(a)** Boxplot for the data showing the significant reduction (p<0.0001) of Evi/Wls expression (blue) in the Vps15 perturbed half. This reduction is converse to the significantly increased intensity signal of Wg in the perturbed half (red; p<0.0001). **(b)** Boxplot of the fluorescence intensity measurements of the extracellular Wg (blue). Extracellular Wg signal in the Vps15 perturbed half is significantly increased (blue; p<0.0001), corresponding to the significant increase (p<0.0001) of the total Wg in the same area (red). Statistical differences between perturbed and unperturbed regions were assessed using the Wilcoxon rank-sum test (wilcox.test) in R. Wt: wildtype; ko: knockout; Exwg: extracellular Wg.

**Supplementary Table S1** | List of all used HD_CFD library sgRNA fly lines. File can be found as .xlsx file.

**Supplementary Table S2** | List of all HD_CFD library sgRNA positive control fly lines used for the CRISPR scree. File can be found as .xlsx file.

**Supplementary Table S3** | List of the initial CRISPR screen hit list. File can be found as .xlsx file.

**Supplementary Table S4** | Sequencing results of the knockout efficiency test with sanger sequencing for sgRNA lines #009 and #83822 crossed to *w-;;act-Gal4,UAS-Cas9.* File can be found as .xlsx file.

**Supplementary Table S5** | Detailed imaging information for the super-resolution data presented in the manuscript. File can be found as .xlsx file.
