## Supplementary material for "Class III PI3K is essential for Wingless secretion and Evi/Wls recycling in *Drosophila*": Table S1

| n | HD_CFDlibrary | Flybase No | gene_name | gene_function |
|---|---|---|---|---|
| 1 | CFDlib00001 | FBgn0032752 | NA | Kinase |
| 2 | CFDlib00003 | FBgn0036428 | Gbs-70E | Phosphatase |
| 3 | CFDlib00004 | FBgn0042711 | Hex-t1 | Phosphatase |
| 4 | CFDlib00005 | FBgn0036266 | NA | Kinase |
| 5 | CFDlib00006 | FBgn0024947 | NTPase | Phosphatase |
| 6 | CFDlib00007 | FBgn0259822 | Ca-beta | Kinase |
| 7 | CFDlib00008 | FBgn0026602 | Ady43A | Kinase |
| 8 | CFDlib00009 | FBgn0260935 | Vps15 | Kinase |
| 9 | CFDlib00011 | FBgn0051183 | NA | Kinase |
| 10 | CFDlib00012 | FBgn0034997 | NA | Phosphatase |
| 11 | CFDlib00014 | FBgn0263855 | BubR1 | Kinase |
| 12 | CFDlib00015 | FBgn0037164 | NA | Phosphatase |
| 13 | CFDlib00016 | FBgn0024326 | Mkk4 | Kinase |
| 14 | CFDlib00017 | FBgn0032820 | fbp | Phosphatase |
| 15 | CFDlib00019 | FBgn0259227 | NA | Phosphatase |
| 16 | CFDlib00020 | FBgn0031397 | CG15385 | Phosphatase |
| 17 | CFDlib00022 | FBgn0260972 | alc | Kinase |
| 18 | CFDlib00023 | FBgn0263980 | \Stacl | Kinase |
| 19 | CFDlib00026 | FBgn0004584 | Rrp1 | Kinase |
| 20 | CFDlib00028 | FBgn0022800 | Cad96Ca | Kinase |
| 21 | CFDlib00029 | FBgn0003256 | rl | Kinase |
| 22 | CFDlib00030 | FBgn0000150 | awd | Kinase |
| 23 | CFDlib00031 | FBgn0265959 | rdgC | Phosphatase |
| 24 | CFDlib00032 | FBgn0037315 | Cerk | Kinase |
| 25 | CFDlib00034 | FBgn0000382 | csw | Phosphatase |
| 26 | CFDlib00035 | FBgn0003502 | Btk29A | Kinase |
| 27 | CFDlib00036 | FBgn0027279 | l(1)G0196 | Kinase |
| 28 | CFDlib00037 | FBgn0031267 | Ipk2 | Kinase |
| 29 | CFDlib00038 | FBgn0002466 | sti | Kinase |
| 30 | CFDlib00039 | FBgn0033773 | Mos | Kinase |
| 31 | CFDlib00040 | FBgn0030018 | slpr | Kinase |
| 32 | CFDlib00041 | FBgn0010015 | CanA1 | Phosphatase |
| 33 | CFDlib00042 | FBgn0034950 | Pask | Kinase |
| 34 | CFDlib00043 | FBgn0039809 | NA | Kinase |
| 35 | CFDlib00044 | FBgn0004373 | fwd | Kinase |
| 36 | CFDlib00046 | FBgn0000464 | Lar | Phosphatase |
| 37 | CFDlib00049 | FBgn0267486 | Ptp36E | Phosphatase |
| 38 | CFDlib00051 | FBgn0032297 | NA | Phosphatase |
| 39 | CFDlib00052 | FBgn0030735 | NA | Phosphatase |
| 40 | CFDlib00053 | FBgn0011725 | twin | Kinase |
| 41 | CFDlib00054 | FBgn0010389 | htl | Kinase |
| 42 | CFDlib00056 | FBgn0000147 | aurA | Kinase |
| 43 | CFDlib00057 | FBgn0260990 | yata | Kinase |
| 44 | CFDlib00059 | FBgn0040079 | pkaap | Kinase |
| 45 | CFDlib00060 | FBgn0259101 | NA | Phosphatase |
| 46 | CFDlib00062 | FBgn0003079 | Raf | Kinase |
| 47 | CFDlib00064 | FBgn0046332 | gskt | Kinase |

| | | | | |
|---|---|---|---|---|
| 48 | CFDlib00067 | FBgn0019686 | lok | Kinase |
| 49 | CFDlib00069 | FBgn0262081 | Csk | Kinase |
| 50 | CFDlib00070 | FBgn0026371 | SAK | Kinase |
| 51 | CFDlib00086 | FBgn0036030 | NA | Kinase |
| 52 | CFDlib00089 | FBgn0264492 | CkIIalpha | Kinase |
| 53 | CFDlib00090 | FBgn0022338 | dnk | Kinase |
| 54 | CFDlib00091 | FBgn0004876 | cdi | Kinase |
| 55 | CFDlib00093 | FBgn0267390 | dop | Kinase |
| 56 | CFDlib00095 | FBgn0010407 | Ror | Kinase |
| 57 | CFDlib00097 | FBgn0034568 | NA | Kinase |
| 58 | CFDlib00098 | FBgn0005666 | bt | Kinase |
| 59 | CFDlib00099 | FBgn0032702 | NA | Phosphatase |
| 60 | CFDlib00102 | FBgn0033791 | Drl-2 | Kinase |
| 61 | CFDlib00103 | FBgn0010441 | pll | Kinase |
| 62 | CFDlib00104 | FBgn0267698 | Pak | Kinase |
| 63 | CFDlib00105 | FBgn0038655 | NA | Phosphatase |
| 64 | CFDlib00107 | FBgn0032677 | CG5790 | Kinase |
| 65 | CFDlib00110 | FBgn0263998 | Ack-like | Kinase |
| 66 | CFDlib00111 | FBgn0260934 | par-1 | Kinase |
| 67 | CFDlib00113 | FBgn0261873 | sdt | Kinase |
| 68 | CFDlib00114 | FBgn0038446 | NA | Kinase |
| 69 | CFDlib00115 | FBgn0020270 | mre11 | Kinase |
| 70 | CFDlib00116 | FBgn0027515 | NA | Phosphatase |
| 71 | CFDlib00117 | FBgn0037327 | PEK | Kinase |
| 72 | CFDlib00118 | FBgn0085385 | bma | Kinase |
| 73 | CFDlib00119 | FBgn0036187 | RIOK1 | Kinase |
| 74 | CFDlib00121 | FBgn0000536 | eas | Kinase |
| 75 | CFDlib00122 | FBgn0030696 | NA | Phosphatase |
| 76 | CFDlib00123 | FBgn0030573 | nmdyn-D6 | Kinase |
| 77 | CFDlib00125 | FBgn0032749 | Phlpp | Phosphatase |
| 78 | CFDlib00126 | FBgn0019990 | Gcn2 | Kinase |
| 79 | CFDlib00128 | FBgn0024329 | Mekk1 | Kinase |
| 80 | CFDlib00129 | FBgn0000274 | Pka-C2 | Kinase |
| 81 | CFDlib00130 | FBgn0027889 | ball | Kinase |
| 82 | CFDlib00131 | FBgn0036337 | Adk2 | Kinase |
| 83 | CFDlib00132 | FBgn0039742 | NA | Phosphatase |
| 84 | CFDlib00133 | FBgn0020440 | Fak | Kinase |
| 85 | CFDlib00134 | FBgn0032779 | Alp12 | Phosphatase |
| 86 | CFDlib00135 | FBgn0000017 | Abl | Kinase |
| 87 | CFDlib00136 | FBgn0052944 | NA | Kinase |
| 88 | CFDlib00137 | FBgn0260945 | Atg1 | Kinase |
| 89 | CFDlib00138 | FBgn0024179 | wit | Kinase |
| 90 | CFDlib00139 | FBgn0005592 | FBgn0285896 | Kinase |
| 91 | CFDlib00140 | FBgn0262617 | Nuak | Kinase |
| 92 | CFDlib00141 | FBgn0011829 | Ret | Kinase |
| 93 | CFDlib00142 | FBgn0003366 | sev | Kinase |
| 94 | CFDlib00143 | FBgn0015380 | drl | Kinase |
| 95 | CFDlib00144 | FBgn0035484 | NA | Kinase |

| | | | | |
|---|---|---|---|---|
| 96 | CFDlib00145 | FBgn0025742 | mtm | Phosphatase |
| 97 | CFDlib00146 | FBgn0243512 | puc | Phosphatase |
| 98 | CFDlib00147 | FBgn0003091 | Pkc53E | Kinase |
| 99 | CFDlib00148 | FBgn0032187 | NA | Kinase |
| 100 | CFDlib00149 | FBgn0035957 | NA | Kinase |
| 101 | CFDlib00150 | FBgn0035421 | nSMase | Phosphatase |
| 102 | CFDlib00151 | FBgn0026063 | KP78b | Kinase |
| 103 | CFDlib00152 | FBgn0032006 | Pvr | Kinase |
| 104 | CFDlib00153 | FBgn0035590 | Prpk | Kinase |
| 105 | CFDlib00154 | FBgn0033673 | NA | Kinase |
| 106 | CFDlib00155 | FBgn0036544 | sff | Kinase |
| 107 | CFDlib00156 | FBgn0026136 | CkIIbeta2 | Kinase |
| 108 | CFDlib00157 | FBgn0004656 | fs(1)h | Kinase |
| 109 | CFDlib00158 | FBgn0004889 | tws | Phosphatase |
| 110 | CFDlib00159 | FBgn0033423 | Alp6 | Phosphatase |
| 111 | CFDlib00160 | FBgn0015277 | Pi3K59F | Kinase |
| 112 | CFDlib00161 | FBgn0035001 | Slik | Kinase |
| 113 | CFDlib00235 | FBgn0031799 | Pez | Phosphatase |
| 114 | CFDlib00236 | FBgn0045980 | niki | Kinase |
| 115 | CFDlib00237 | FBgn0036161 | NA | Kinase |
| 116 | CFDlib00238 | FBgn0261984 | Ire1 | Kinase |
| 117 | CFDlib00239 | FBgn0031952 | cdc14 | Phosphatase |
| 118 | CFDlib00240 | FBgn0052487 | NA | Phosphatase |
| 119 | CFDlib00242 | FBgn0000711 | flw | Phosphatase |
| 120 | CFDlib00245 | FBgn0005536 | Mbs | Phosphatase |
| 121 | CFDlib00246 | FBgn0000405 | CycB | Kinase |
| 122 | CFDlib00247 | FBgn0031044 | MKP-4 | Phosphatase |
| 123 | CFDlib00248 | FBgn0086657 | IKKepsilon | Kinase |
| 124 | CFDlib00249 | FBgn0034644 | CG10082 | Kinase |
| 125 | CFDlib00250 | FBgn0000826 | png | Kinase |
| 126 | CFDlib00251 | FBgn0016930 | Dyrk2 | Kinase |
| 127 | CFDlib00252 | FBgn0032811 | Pmvk | Kinase |
| 128 | CFDlib00253 | FBgn0034691 | Synj | Phosphatase |
| 129 | CFDlib00254 | FBgn0000063 | Mps1 | Kinase |
| 130 | CFDlib00257 | FBgn0015279 | Pi3K92E | Kinase |
| 131 | CFDlib00258 | FBgn0024846 | p38b | Kinase |
| 132 | CFDlib00259 | FBgn0027506 | EDTP | Phosphatase |
| 133 | CFDlib00260 | FBgn0000116 | Argk | Kinase |
| 134 | CFDlib00261 | FBgn0011300 | babo | Kinase |
| 135 | CFDlib00263 | FBgn0036142 | Adck5 | Kinase |
| 136 | CFDlib00264 | FBgn0000442 | Pkg21D | Kinase |
| 137 | CFDlib00265 | FBgn0023169 | AMPKalpha | Kinase |
| 138 | CFDlib00266 | FBgn0259168 | mnb | Kinase |
| 139 | CFDlib00267 | FBgn0004369 | Ptp99A | Phosphatase |
| 140 | CFDlib00268 | FBgn0030245 | NA | Phosphatase |
| 141 | CFDlib00270 | FBgn0260399 | gwl | Kinase |
| 142 | CFDlib00273 | FBgn0038463 | NA | Kinase |
| 143 | CFDlib00274 | FBgn0036552 | NA | Phosphatase |

| | | | | |
|---|---|---|---|---|
| 144 | CFDlib00275 | FBgn0034137 | NA | Kinase |
| 145 | CFDlib00277 | FBgn0032650 | CG7094 | Kinase |
| 146 | CFDlib00278 | FBgn0022029 | l(2)k01209 | Kinase |
| 147 | CFDlib00279 | FBgn0085386 | NA | Kinase |
| 148 | CFDlib00280 | FBgn0035945 | NA | Phosphatase |
| 149 | CFDlib00282 | FBgn0015618 | Cdk8 | Kinase |
| 150 | CFDlib00284 | FBgn0027507 | NA | Kinase |
| 151 | CFDlib00285 | FBgn0028497 | Mtmr6 | Phosphatase |
| 152 | CFDlib00286 | FBgn0028427 | Ilk | Kinase |
| 153 | CFDlib00287 | FBgn0031995 | NA | Kinase |
| 154 | CFDlib00288 | FBgn0052484 | Sk2 | Kinase |
| 155 | CFDlib00291 | FBgn0021796 | Tor | Kinase |
| 156 | CFDlib00292 | FBgn0040298 | Myt1 | Kinase |
| 157 | CFDlib00294 | FBgn0027497 | Madm | Kinase |
| 158 | CFDlib00296 | FBgn0039796 | NA | Kinase |
| 159 | CFDlib00297 | FBgn0035039 | Adck1 | Kinase |
| 160 | CFDlib00298 | FBgn0038744 | NA | Phosphatase |
| 161 | CFDlib00299 | FBgn0023129 | aay | Phosphatase |
| 162 | CFDlib00300 | FBgn0030556 | mRNA-cap | Phosphatase |
| 163 | CFDlib00301 | FBgn0036028 | NA | Phosphatase |
| 164 | CFDlib00302 | FBgn0031463 | G6P | Phosphatase |
| 165 | CFDlib00303 | FBgn0040394 | NA | Kinase |
| 166 | CFDlib00304 | FBgn0037218 | aux | Kinase |
| 167 | CFDlib00305 | FBgn0003744 | trc | Kinase |
| 168 | CFDlib00306 | FBgn0261549 | rdgA | Kinase |
| 169 | CFDlib00307 | FBgn0031730 | NA | Kinase |
| 170 | CFDlib00308 | FBgn0050103 | NA | Phosphatase |
| 171 | CFDlib00309 | FBgn0010316 | dap | Kinase |
| 172 | CFDlib00310 | FBgn0023508 | Ocrl | Phosphatase |
| 173 | CFDlib00311 | FBgn0036844 | Mkp3 | Phosphatase |
| 174 | CFDlib00313 | FBgn0028741 | fab1 | Kinase |
| 175 | CFDlib00315 | FBgn0036551 | NA | Phosphatase |
| 176 | CFDlib00316 | FBgn0030208 | PPP4R2r | Phosphatase |
| 177 | CFDlib00317 | FBgn0003169 | put | Kinase |
| 178 | CFDlib00318 | FBgn0267350 | Pi4KIIIalpha | Kinase |
| 179 | CFDlib00399 | FBgn0264357 | SNF4Agamma | Kinase |
| 180 | CFDlib00400 | FBgn0036760 | NA | Phosphatase |
| 181 | CFDlib00401 | FBgn0261456 | hpo | Kinase |
| 182 | CFDlib00404 | FBgn0046685 | Wsck | Kinase |
| 183 | CFDlib00405 | FBgn0265998 | Doa | Kinase |
| 184 | CFDlib00406 | FBgn0026061 | Mipp1 | Phosphatase |
| 185 | CFDlib00516 | FBgn0263968 | nonC | Kinase |
| 186 | CFDlib00517 | FBgn0028360 | Cdc7 | Kinase |
| 187 | CFDlib00519 | FBgn0003733 | tor | Kinase |
| 188 | CFDlib00520 | FBgn0033322 | NA | Phosphatase |
| 189 | CFDlib00521 | FBgn0004784 | inaC | Kinase |
| 190 | CFDlib00524 | FBgn0032147 | IP3K1 | Kinase |
| 191 | CFDlib00525 | FBgn0050295 | Ipk1 | Kinase |

| | | | | |
|---|---|---|---|---|
| 192 | CFDlib00711 | FBgn0038167 | Lkb1 | Kinase |
| 193 | CFDlib00712 | FBgn0015772 | Nak | Kinase |
| 194 | CFDlib00713 | FBgn0037167 | NA | Phosphatase |
| 195 | CFDlib00714 | FBgn0029736 | NA | Kinase |
| 196 | CFDlib00715 | FBgn0036876 | NA | Phosphatase |
| 197 | CFDlib00716 | FBgn0030697 | NA | Kinase |
| 198 | CFDlib00717 | FBgn0038630 | NA | Kinase |
| 199 | CFDlib00718 | FBgn0260750 | Mulk | Kinase |
| 200 | CFDlib00720 | FBgn0036896 | wnd | Kinase |
| 201 | CFDlib00721 | FBgn0020391 | Nrk | Kinase |
| 202 | CFDlib00722 | FBgn0039306 | RIOK2 | Kinase |
| 203 | CFDlib00723 | FBgn0029958 | Pdp | Phosphatase |
| 204 | CFDlib00724 | FBgn0026181 | Rok | Kinase |
| 205 | CFDlib00727 | FBgn0016672 | Ipp | Phosphatase |
| 206 | CFDlib00729 | FBgn0061359 | NA | Kinase |
| 207 | CFDlib00732 | FBgn0036877 | NA | Phosphatase |
| 208 | CFDlib00734 | FBgn0036273 | INPP5E | Phosphatase |
| 209 | CFDlib00735 | FBgn0034710 | Alp7 | Phosphatase |
| 210 | CFDlib00736 | FBgn0052549 | NA | Phosphatase |
| 211 | CFDlib00738 | FBgn0036553 | NA | Phosphatase |
| 212 | CFDlib00739 | FBgn0037093 | Cdk12 | Kinase |
| 213 | CFDlib00744 | FBgn0023083 | fray | Kinase |
| 214 | CFDlib00747 | FBgn0037063 | NA | Phosphatase |
| 215 | CFDlib00748 | FBgn0035619 | Alp10 | Phosphatase |
| 216 | CFDlib00749 | FBgn0263395 | hppy | Kinase |
| 217 | CFDlib00750 | FBgn0025640 | NA | Kinase |
| 218 | CFDlib00751 | FBgn0250906 | Pgk | Kinase |
| 219 | CFDlib00754 | FBgn0016984 | sktl | Kinase |
| 220 | CFDlib00756 | FBgn0017581 | Lk6 | Kinase |
| 221 | CFDlib00758 | FBgn0261278 | grp | Kinase |
| 222 | CFDlib00760 | FBgn0014007 | Ptp69D | Phosphatase |
| 223 | CFDlib00773 | FBgn0002938 | ninaC | Kinase |
| 224 | CFDlib00776 | FBgn0031030 | Tao | Kinase |
| 225 | CFDlib00779 | FBgn0035426 | NA | Phosphatase |
| 226 | CFDlib00782 | FBgn0035142 | Hipk | Kinase |
| 227 | CFDlib00783 | FBgn0003132 | Pp1-13C | Phosphatase |
| 228 | CFDlib00800 | FBgn0022768 | Pp2C1 | Phosphatase |
| 229 | CFDlib00801 | FBgn0036550 | NA | Phosphatase |
| 230 | CFDlib00806 | FBgn0014006 | Ask1 | Kinase |
| 231 | CFDlib00807 | FBgn0025743 | mbt | Kinase |
| 232 | CFDlib00808 | FBgn0266465 | GckIII | Kinase |
| 233 | CFDlib00809 | FBgn0028484 | Ack | Kinase |
| 234 | CFDlib00811 | FBgn0015765 | p38a | Kinase |
| 235 | CFDlib00812 | FBgn0016641 | PTP-ER | Phosphatase |
| 236 | CFDlib00814 | FBgn0029157 | ssh | Phosphatase |
| 237 | CFDlib00828 | FBgn0039015 | Takl2 | Kinase |
| 238 | CFDlib00830 | FBgn0031696 | Bub1 | Kinase |
| 239 | CFDlib00832 | FBgn0003731 | Egfr | Kinase |

| | | | | |
|---|---|---|---|---|
| 240 | CFDlib01038 | FBgn0023097 | bon | Kinase |
| 241 | CFDlib01345 | FBgn0024222 | IKKbeta | Kinase |
| 242 | CFDlib01346 | FBgn0016696 | Pitslre | Kinase |
| 243 | CFDlib01347 | FBgn0004839 | otk | Kinase |
| 244 | CFDlib01348 | FBgn0024227 | aurB | Kinase |
| 245 | CFDlib01349 | FBgn0015402 | ksr | Kinase |
| 246 | CFDlib01350 | FBgn0002413 | dco | Kinase |
| 247 | CFDlib01352 | FBgn0286813 | SRPK | Kinase |
| 248 | CFDlib01353 | FBgn0010355 | Taf1 | Kinase |
| 249 | CFDlib01354 | FBgn0026064 | KP78a | Kinase |
| 250 | CFDlib01355 | FBgn0011739 | wts | Kinase |
| 251 | CFDlib01356 | FBgn0010269 | Dsor1 | Kinase |
| 252 | CFDlib01357 | FBgn0023081 | gek | Kinase |
| 253 | CFDlib01358 | FBgn0025936 | Eph | Kinase |
| 254 | CFDlib01359 | FBgn0013987 | MAPk-Ak2 | Kinase |
| 255 | CFDlib01360 | FBgn0020412 | JIL-1 | Kinase |
| 256 | CFDlib01361 | FBgn0001079 | fu | Kinase |
| 257 | CFDlib01362 | FBgn0005640 | Eip63E | Kinase |
| 258 | CFDlib01363 | FBgn0015024 | CkIalpha | Kinase |
| 259 | CFDlib01364 | FBgn0025702 | Srpk79D | Kinase |
| 260 | CFDlib01365 | FBgn0024245 | dnt | Kinase |
| 261 | CFDlib01366 | FBgn0004864 | hop | Kinase |
| 262 | CFDlib01367 | FBgn0010379 | Akt1 | Kinase |
| 263 | CFDlib01368 | FBgn0027101 | Dyrk3 | Kinase |
| 264 | CFDlib01369 | FBgn0013762 | Cdk5 | Kinase |
| 265 | CFDlib01370 | FBgn0000229 | bsk | Kinase |
| 266 | CFDlib01371 | FBgn0020621 | Pkn | Kinase |
| 267 | CFDlib01372 | FBgn0019949 | Cdk9 | Kinase |
| 268 | CFDlib01374 | FBgn0004367 | mei-41 | Kinase |
| 269 | CFDlib01375 | FBgn0010909 | msn | Kinase |
| 270 | CFDlib01376 | FBgn0027504 | NA | Kinase |
| 271 | CFDlib01377 | FBgn0010303 | hep | Kinase |
| 272 | CFDlib01378 | FBgn0011737 | Wee1 | Kinase |
| 273 | CFDlib01389 | FBgn0040056 | NA | Kinase |
| 274 | CFDlib01393 | FBgn0015295 | Shark | Kinase |
| 275 | CFDlib01399 | FBgn0031643 | NA | Kinase |
| 276 | CFDlib01400 | FBgn0265045 | Strn-Mlck | Kinase |
| 277 | CFDlib01401 | FBgn0030384 | NA | Kinase |
| 278 | CFDlib01402 | FBgn0031441 | NA | Kinase |
| 279 | CFDlib01403 | FBgn0037098 | Wnk | Kinase |
| 280 | CFDlib01404 | FBgn0039908 | Asator | Kinase |
| 281 | CFDlib01405 | FBgn0044826 | Pak3 | Kinase |
| 282 | CFDlib01406 | FBgn0250823 | gish | Kinase |
| 283 | CFDlib01407 | FBgn0053554 | Nipped-A | Kinase |
| 284 | CFDlib01410 | FBgn0000723 | FER | Kinase |
| 285 | CFDlib01411 | FBgn0020386 | Pdk1 | Kinase |
| 286 | CFDlib01412 | FBgn0026323 | Tak1 | Kinase |
| 287 | CFDlib01413 | FBgn0262733 | Src64B | Kinase |

| | | | | |
|---|---|---|---|---|
| 288 | CFDlib01414 | FBgn0013435 | Cdc2rk | Kinase |
| 289 | CFDlib01418 | FBgn0039083 | NA | Kinase |
| 290 | CFDlib01419 | FBgn0053519 | Unc-89 | Kinase |
| 291 | CFDlib01420 | FBgn0260798 | Gprk1 | Kinase |
| 292 | CFDlib01421 | FBgn0031299 | NA | Kinase |
| 293 | CFDlib01422 | FBgn0000658 | fj | Kinase |
| 294 | CFDlib01423 | FBgn0038816 | Lrrk | Kinase |
| 295 | CFDlib01424 | FBgn0262866 | S6kII | Kinase |
| 296 | CFDlib01425 | FBgn0283473 | S6KL | Kinase |
| 297 | CFDlib01426 | FBgn0283657 | Tlk | Kinase |
| 298 | CFDlib01428 | FBgn0285896 | btl | Kinase |
| 299 | CFDlib01429 | FBgn0053531 | Ddr | Kinase |
| 300 | CFDlib01431 | FBgn0033915 | NA | Kinase |
| 301 | CFDlib01433 | FBgn0031855 | meng | Kinase |
| 302 | CFDlib01435 | FBgn0263237 | Cdk7 | Kinase |
| 303 | CFDlib01436 | FBgn0264087 | Slob | Kinase |
| 304 | CFDlib01437 | FBgn0040505 | Alk | Kinase |
| 305 | CFDlib01438 | FBgn0035397 | PAN3 | Kinase |
| 306 | CFDlib01439 | FBgn0046689 | Takl1 | Kinase |
| 307 | CFDlib01440 | FBgn0261854 | aPKC | Kinase |
| 308 | CFDlib01442 | FBgn0004106 | Cdk1 | Kinase |
| 309 | CFDlib01443 | FBgn0283472 | S6k | Kinase |
| 310 | CFDlib01445 | FBgn0046706 | Haspin | Kinase |
| 311 | CFDlib01446 | FBgn0283499 | InR | Kinase |
| 312 | CFDlib01447 | FBgn0264959 | Src42A | Kinase |
| 313 | CFDlib01449 | FBgn0045035 | tefu | Kinase |
| 314 | CFDlib01450 | FBgn0011754 | PhKgamma | Kinase |
| 315 | CFDlib01455 | FBgn0259678 | sqa | Kinase |
| 316 | CFDlib01456 | FBgn0003716 | tkv | Kinase |
| 317 | CFDlib01473 | FBgn0032424 | NA | Kinase |
| 318 | CFDlib01483 | FBgn0032083 | NA | Kinase |
| 319 | CFDlib01504 | FBgn0034085 | Ptp52F | Phosphatase |
| 320 | CFDlib01514 | FBgn0037578 | PNKP | Phosphatase |
| 321 | CFDlib01529 | FBgn0020389 | Papss | Kinase |
| 322 | CFDlib01537 | FBgn0004107 | Cdk2 | Kinase |
| 323 | CFDlib01577 | FBgn0024734 | PRL-1 | Phosphatase |
| 324 | CFDlib01595 | FBgn0030661 | Alp11 | Phosphatase |
| 325 | CFDlib01600 | FBgn0003525 | stg | Phosphatase |
| 326 | CFDlib01618 | FBgn0028410 | Pk34A | Kinase |
| 327 | CFDlib01621 | FBgn0031907 | NA | Phosphatase |
| 328 | CFDlib01646 | FBgn0028833 | Cmpk | Kinase |
| 329 | CFDlib01653 | FBgn0003317 | sax | Kinase |
| 330 | CFDlib01657 | FBgn0002673 | twe | Phosphatase |
| 331 | CFDlib01678 | FBgn0023177 | Pp4-19C | Phosphatase |
| 332 | CFDlib01682 | FBgn0031784 | NA | Kinase |
| 333 | CFDlib01724 | FBgn0037786 | Alp13 | Phosphatase |
| 334 | CFDlib01746 | FBgn0025802 | Sbf | Phosphatase |
| 335 | CFDlib01747 | FBgn0000032 | Acph-1 | Phosphatase |

| | | | | |
|---|---|---|---|---|
| 336 | CFDlib01752 | FBgn0004103 | Pp1-87B | Phosphatase |
| 337 | CFDlib01779 | FBgn0034789 | PIP5K59B | Kinase |
| 338 | CFDlib01782 | FBgn0030465 | NA | Phosphatase |
| 339 | CFDlib01795 | FBgn0020930 | Dgkepsilon | Kinase |
| 340 | CFDlib01810 | FBgn0017558 | Pdk | Kinase |
| 341 | CFDlib01812 | FBgn0030300 | Sk1 | Kinase |
| 342 | CFDlib01832 | FBgn0035425 | NA | Phosphatase |
| 343 | CFDlib01834 | FBgn0036942 | NA | Kinase |
| 344 | CFDlib01842 | FBgn0014930 | NA | Kinase |
| 345 | CFDlib01843 | FBgn0036369 | NA | Phosphatase |
| 346 | CFDlib01874 | FBgn0037166 | NA | Phosphatase |
| 347 | CFDlib01876 | FBgn0037491 | NA | Kinase |
| 348 | CFDlib01896 | FBgn0062449 | NA | Phosphatase |
| 349 | CFDlib01905 | FBgn0051140 | NA | Kinase |
| 350 | CFDlib01953 | FBgn0028341 | Ptpmeg2 | Phosphatase |
| 351 | CFDlib01956 | FBgn0011205 | fbl | Kinase |
| 352 | CFDlib01964 | FBgn0000489 | Pka-C3 | Kinase |
| 353 | CFDlib01967 | FBgn0036160 | NA | Kinase |
| 354 | CFDlib01978 | FBgn0034299 | NA | Kinase |
| 355 | CFDlib01983 | FBgn0011826 | Pp2B-14D | Phosphatase |
| 356 | CFDlib01985 | FBgn0037339 | Pi4KIIalpha | Kinase |
| 357 | CFDlib01993 | FBgn0004177 | mts | Phosphatase |
| 358 | CFDlib02000 | FBgn0035228 | NA | Phosphatase |
| 359 | CFDlib02020 | FBgn0036448 | mop | Phosphatase |
| 360 | CFDlib02023 | FBgn0036875 | NA | Phosphatase |
| 361 | CFDlib02026 | FBgn0000721 | for | Kinase |
| 362 | CFDlib02044 | FBgn0003134 | Pp1alpha-96A | Phosphatase |
| 363 | CFDlib02057 | FBgn0003140 | PpY-55A | Phosphatase |
| 364 | CFDlib02061 | FBgn0046692 | Stlk | Kinase |
| 365 | CFDlib02080 | FBgn0029970 | Nek2 | Kinase |
| 366 | CFDlib02094 | FBgn0015278 | Pi3K68D | Kinase |
| 367 | CFDlib02098 | FBgn0025573 | PpN58A | Phosphatase |
| 368 | CFDlib02100 | FBgn0261524 | lic | Kinase |
| 369 | CFDlib02119 | FBgn0004368 | Ptp4E | Phosphatase |
| 370 | CFDlib02139 | FBgn0259166 | NA | Phosphatase |
| 371 | CFDlib02148 | FBgn0029067 | Dd | Phosphatase |
| 372 | CFDlib02160 | FBgn0038588 | NA | Kinase |
| 373 | CFDlib02163 | FBgn0005777 | PpD3 | Phosphatase |
| 374 | CFDlib02167 | FBgn0263398 | Uck | Kinase |
| 375 | CFDlib02172 | FBgn0034712 | Alp8 | Phosphatase |
| 376 | CFDlib02186 | FBgn0038912 | NA | Phosphatase |
| 377 | CFDlib02212 | FBgn0005779 | PpD6 | Phosphatase |
| 378 | CFDlib02235 | FBgn0028978 | trbl | Kinase |
| 379 | CFDlib02237 | FBgn0261387 | NA | Kinase |
| 380 | CFDlib02239 | FBgn0029949 | NA | Phosphatase |
| 381 | CFDlib02247 | FBgn0025592 | Gk1 | Kinase |
| 382 | CFDlib02276 | FBgn0031451 | NA | Kinase |
| 383 | CFDlib02285 | FBgn0026060 | Mipp2 | Phosphatase |

| | | | | |
|---|---|---|---|---|
| 384 | CFDlib02287 | FBgn0041087 | wun2 | Phosphatase |
| 385 | CFDlib02307 | FBgn0029891 | Pink1 | Kinase |
| 386 | CFDlib02342 | FBgn0016126 | CaMKI | Kinase |
| 387 | CFDlib02363 | FBgn0052666 | Drak | Kinase |
| 388 | CFDlib02366 | FBgn0027621 | Pfrx | Phosphatase |
| 389 | CFDlib02376 | FBgn0042094 | Ak3 | Kinase |
| 390 | CFDlib02390 | FBgn0001624 | dlg1 | Kinase |
| 391 | CFDlib02392 | FBgn0263199 | Galk | Kinase |
| 392 | CFDlib02395 | FBgn0039698 | NA | Phosphatase |
| 393 | CFDlib02406 | FBgn0038902 | NA | Kinase |
| 394 | CFDlib02435 | FBgn0040077 | primo-1 | Phosphatase |
| 395 | CFDlib02473 | FBgn0051145 | CG31145 | Kinase |
| 396 | CFDlib02491 | FBgn0052703 | Erk7 | Kinase |
| 397 | CFDlib02500 | FBgn0003139 | PpV | Phosphatase |
| 398 | CFDlib02503 | FBgn0259178 | 5PtaseI | Phosphatase |
| 399 | CFDlib02520 | FBgn0038603 | PKD | Kinase |
| 400 | CFDlib02523 | FBgn0037325 | NA | Kinase |
| 401 | CFDlib02547 | FBgn0033021 | NA | Phosphatase |
| 402 | CFDlib02574 | FBgn0022709 | Ak1 | Kinase |
| 403 | CFDlib02592 | FBgn0030976 | NA | Phosphatase |
| 404 | CFDlib00905 | FBgn0004360 | Wnt2 | pc |
| 405 | CFDlib01259 | FBgn0000119 | arr | pc |
| 406 | CFDlib01623 | FBgn0031902 | Wnt6 | pc |
| 407 | CFDlib01770 | FBgn0010194 | Wnt5 | pc |
| 408 | CFDlib01931 | FBgn0250823 | gish | pc |
| 409 | CFDlib01954 | FBgn0036141 | wls | pc |
| 410 | CFDlib02056 | FBgn0010194 | Wnt5 | pc |
| 411 | CFDlib02175 | FBgn0003134 | Pp1alpha-96A | Phosphatase, pc |
