## Supplementary material for "Class III PI3K is essential for Wingless secretion and Evi/Wls recycling in *Drosophila*": Table S2

| n | HD_CFD_library | Flybase No. | Gene name | positive control | gene_function | Source | Source 2 | Phenotype | Effect on (after Swarup et al., 2015) | Pheno |
|---|---|---|---|---|---|---|---|---|---|---|
| 1 | CFDlib00030 | FBgn0000150 | awd | pc | Kinase | Swarup et al., | Gross et al., | yes | sens, dll, wg, ct | |
| 2 | CFDlib00089 | FBgn0264492 | CkIIalpha | pc | Kinase | Swarup et al., | Gross et al., | yes | sens, dll, wg, ct | |
| 3 | CFDlib00095 | FBgn0010407 | Ror | pc | other | known | | yes | | |
| 4 | CFDlib00111 | FBgn0260934 | par-1 | pc | Kinase | Swarup et al., | | yes | sens, dll, wg, ct | |
| 5 | CFDlib00126 | FBgn0019990 | Gcn2 | pc | Kinase | Swarup et al., | | yes | sens, dll, wg, ct | |
| 6 | CFDlib00145 | FBgn0025742 | mtm | pc | Kinase | Swarup et al., | Chaudhary et al | yes | sens, dll, wg, ct | |
| 7 | CFDlib00156 | FBgn0026136 | CkIIbeta2 | pc | Kinase | Swarup et al., | | yes | sens, dll, wg, ct | |
| 8 | CFDlib00235 | FBgn0031799 | Pez | pc | Phosphatase | Swarup et al., | | yes | sens, dll, wg, ct | |
| 9 | CFDlib00239 | FBgn0031952 | cdc14 | pc | Phosphatase | known | | yes | | |
| 10 | CFDlib00242 | FBgn0000711 | flw | pc | Phosphatase | Swarup et al., | | yes | sens, dll, wg, ct | |
| 11 | CFDlib00247 | FBgn0031044 | MKP-4 | pc | Phosphatase | Swarup et al., | | yes | sens, dll, wg, ct | |
| 12 | CFDlib00253 | FBgn0034691 | Synj | pc | Phosphatase | Swarup et al., | | yes | sens, dll, wg, ct | |
| 13 | CFDlib00294 | FBgn0027497 | Madm | pc | other | Chaudhary et al., | | lethal | | |
| 14 | CFDlib00300 | FBgn0030556 | mRNA-cap | pc | Phosphatase | Swarup et al., | | yes | sens, dll, wg, ct | |
| 15 | CFDlib00304 | FBgn0037218 | aux | pc | Kinase | Swarup et al., | Chaudhary et al | yes | sens, dll, wg, ct | |
| 16 | CFDlib00316 | FBgn0030208 | PPP4R2r | pc | Phosphatase | Swarup et al., | | yes | sens, dll, wg, ct | |
| 17 | CFDlib00524 | FBgn0032147 | IP3K1 | pc | Kinase | Swarup et al., | | yes | sens, dll, wg, ct | |
| 18 | CFDlib00739 | FBgn0037093 | Cdk12 | pc | Kinase | Swarup et al., | | yes | sens, dll, wg, ct | |
| 19 | CFDlib00782 | FBgn0035142 | Hipk | pc | Kinase | Swarup et al., | | yes | sens, dll, wg, ct | |
| 20 | CFDlib00783 | FBgn0003132 | Pp1-13C | pc | Phosphatase | Swarup et al., | | yes | sens, dll, wg, ct | |
| 21 | CFDlib00800 | FBgn0022768 | Pp2C1 | pc | Phosphatase | Swarup et al., | | yes | sens, dll, wg, ct | |
| 22 | CFDlib00830 | FBgn0031696 | Bub1 | pc | Kinase | Swarup et al., | | yes | sens, dll, wg, ct | |
| 23 | CFDlib01346 | FBgn0016696 | Pitslre | pc | Kinase | Swarup et al., | | yes | sens, dll, wg, ct | |
| 24 | CFDlib01347 | FBgn0004839 | otk | pc | Kinase | Swarup et al., | | yes | sens, dll, wg, ct | |
| 25 | CFDlib01362 | FBgn0005640 | Eip63E | pc | Kinase | Swarup et al., | | yes | sens, dll, wg, ct | |
| 26 | CFDlib01363 | FBgn0015024 | CkIalpha | pc | Kinase | Swarup et al., | | yes | sens, dll, wg, ct | |
| 27 | CFDlib01372 | FBgn0019949 | Cdk9 | pc | Kinase | Swarup et al., | | yes | sens, dll, wg, ct | |
| 28 | CFDlib01406 | FBgn0250823 | gish | pc | Kinase | Swarup et al., | Gross et al., | yes | sens, dll, wg, ct | |
| 29 | CFDlib01407 | FBgn0053554 | Nipped-A | pc | Kinase | Swarup et al., | | yes | sens, dll, wg, ct | |
| 30 | CFDlib01414 | FBgn0013435 | Cdc2rk | pc | Kinase | Swarup et al., | | yes | sens, dll, wg, ct | |
| 31 | CFDlib01420 | FBgn0260798 | Gprk1 | pc | Kinase | Swarup et al., | | yes | sens, dll, wg, ct | |
| 32 | CFDlib01426 | FBgn0283657 | Tlk | pc | Kinase | Swarup et al., | | yes | sens, dll, wg, ct | |
| 33 | CFDlib01435 | FBgn0263237 | Cdk7 | pc | Kinase | known | | yes | | |
| 34 | CFDlib01600 | FBgn0003525 | stg | pc | Phosphatase | Swarup et al., | | yes | sens, dll, wg, ct | |
| 35 | CFDlib01678 | FBgn0023177 | Pp4-19C | pc | Phosphatase | Swarup et al., | | yes | sens, dll, wg, ct | |
| 36 | CFDlib01752 | FBgn0004103 | Pp1-87B | pc | Phosphatase | Swarup et al., | | yes | sens, dll, wg, ct | |
| 37 | CFDlib01931 | FBgn0250823 | gish | pc | Kinase | Swarup et al., | | yes | sens, dll, wg, ct | |
| 38 | CFDlib01953 | FBgn0028341 | l(1)G0232 | pc | other | Swarup et al., | Chaudhary et al | lethal | sens, dll, wg, ct | |
| 39 | CFDlib01954 | FBgn0036141 | wls | pc | other | Chaudhary et al | known | yes | | |
| 40 | CFDlib01993 | FBgn0004177 | mts | pc | Phosphatase | Swarup et al., | | yes | sens, dll, wg, ct | |
| 41 | CFDlib02020 | FBgn0036448 | mop | pc | Phosphatase | Swarup et al., | Chaudhary et al | yes | sens, dll, wg | |
| 42 | CFDlib02026 | FBgn0000721 | for | pc | Kinase | Swarup et al., | | yes | sens, dll, wg, ct | |
| 43 | CFDlib02044 | FBgn0003134 | Pp1alpha-96A | pc | Phosphatase | Swarup et al., | Gross et al., | yes | sens, dll, wg, ct | |
| 44 | CFDlib02163 | FBgn0005777 | PpD3 | pc | Kinase | Swarup et al., | | yes | sens, dll, wg, ct | |
| 45 | CFDlib02175 | FBgn0003134 | Pp1alpha-96A | pc | Phosphatase | Swarup et al., | Gross et al., | yes | sens, dll, wg, ct | |
| 46 | CFDlib02342 | FBgn0016126 | CaMKI | pc | Kinase | known | | yes | | |
| 47 | CFDlib02363 | FBgn0052666 | Drak | pc | Kinase | Swarup et al., | | no | sens, dll, wg, ct | |
| 48 | CFDlib02390 | FBgn0001624 | dlg1 | pc | Kinase | Swarup et al., | | yes | sens, dll, wg, ct | |
| 49 | CFDlib02391 | FBgn0004177 | mts | pc | Kinase | Swarup et al., | | yes | sens, dll, wg, ct | |
| 50 | CFDlib02435 | FBgn0040077 | primo-1 | pc | Phosphatase | Swarup et al., | | yes | sens, dll, wg, ct | |
| 51 | CFDlib02500 | FBgn0003139 | PpV | pc | Phosphatase | Swarup et al., | | yes | sens, dll, wg, ct | |
