## Supplementary material for "Class III PI3K is essential for Wingless secretion and Evi/Wls recycling in *Drosophila*": Table S3

| Stock | Flybase No. | Gene name | Kinase or Phosphatase | Position | Wg Phneotype | Level of penetrance |
|---|---|---|---|---|---|---|
| CFDlib00009 | FBgn0260935 | Vps15 | Kinase | 3R:9,240,757..9,245,869 [-] | strong increase | high |
| CFDlib00016 | FBgn0024326 | Mkk4 | Kinase | 3R:8,646,172..8,650,796 [-] | mild reduction | medium |
| CFDlib00828 | FBgn0039015 | Takl2 | Kinase | 3R:22,732,356..22,733,851 [-] | mild reduction | medium |
| CFDlib02307 | FBgn0029891 | Pink1 | Kinase | X:6,683,798..6,687,302 [+] | mild increase | low |
