## Supplementary material for "Class III PI3K is essential for Wingless secretion and Evi/Wls recycling in *Drosophila*": Table S4

**Sanger Sequnecing results of sgRNA009 and analysis on ICE**

| Label | ICE | KO-Score | KI-Score | ICE d | R Squared | Mean Discord Before | Mean Discord After | Guide Sequences | Control Sample Quality Score | Edit Sample Quality Score | Indels |
|---|---|---|---|---|---|---|---|---|---|---|---|
| KIN173 | 76 | 40 | | 93 | 0.82 | 0.017896423 | 0.52622283 | ACAAGGCAGCCTACATAATG | 58 | 58 | {'0': 6.0, '-1': 8.0, '-3': 25.0, '-5': 6.0, '-7': 7.0, '-8': 4.0, '-9': 10.0, '-10': 11.0, '-11': 4.0, '-12': 1.0} |
| KIN180 | 69 | 39 | | 88 | 0.79 | 0.016595225 | 0.55092836 | ACAAGGCAGCCTACATAATG | 58 | 58 | {'0': 10.0, '-1': 11.0, '-3': 26.0, '-5': 1.0, '-7': 5.0, '-8': 5.0, '-9': 1.0, '-10': 8.0, '-11': 7.0, '-12': 3.0, '-13': 2.0} |
| KIN177 | 79 | 41 | | 100 | 0.79 | 0.013773084 | 0.677664699 | ACAAGGCAGCCTACATAATG | 58 | 58 | {'1': 1.0, '-1': 5.0, '-3': 22.0, '-5': 5.0, '-7': 6.0, '-8': 5.0, '-9': 8.0, '-10': 10.0, '-11': 9.0, '-12': 8.0} |
| KIN176 | 79 | 43 | | 100 | 0.79 | 0.006532148 | 0.710580424 | ACAAGGCAGCCTACATAATG | 58 | 58 | {'-1': 8.0, '-3': 19.0, '-5': 2.0, '-7': 9.0, '-8': 5.0, '-9': 15.0, '-10': 5.0, '-11': 14.0, '-12': 2.0} |
| KIN179 | 78 | 47 | | 100 | 0.78 | 0.014785901 | 0.649694538 | ACAAGGCAGCCTACATAATG | 58 | 61 | {'-1': 5.0, '-3': 15.0, '-5': 6.0, '-7': 11.0, '-8': 5.0, '-9': 10.0, '-10': 9.0, '-11': 10.0, '-12': 3.0, '-15': 3.0, '-16': 1.0} |
| KIN174 | 79 | 56 | | 100 | 0.79 | 0.016055497 | 0.615054633 | ACAAGGCAGCCTACATAATG | 58 | 61 | {'-1': 13.0, '-3': 15.0, '-5': 3.0, '-7': 10.0, '-8': 8.0, '-9': 2.0, '-10': 8.0, '-11': 14.0, '-12': 6.0} |
| KIN181 | 79 | 35 | | 96 | 0.84 | 0.014595707 | 0.563008444 | ACAAGGCAGCCTACATAATG | 58 | 61 | {'0': 5.0, '-1': 7.0, '-3': 36.0, '-5': 2.0, '-7': 8.0, '-8': 4.0, '-9': 5.0, '-10': 6.0, '-11': 8.0, '-12': 3.0} |
| KIN175 | 73 | 32 | | 89 | 0.83 | 0.0141558 | 0.510801023 | ACAAGGCAGCCTACATAATG | 58 | 61 | {'0': 10.0, '-1': 4.0, '-3': 33.0, '-5': 6.0, '-7': 6.0, '-9': 7.0, '-10': 8.0, '-11': 8.0, '-12': 1.0} |
| KIN178 | 79 | 44 | | 100 | 0.79 | 0.014885206 | 0.69004961 | ACAAGGCAGCCTACATAATG | 58 | 61 | {'1': 7.0, '-1': 7.0, '-3': 22.0, '-5': 4.0, '-7': 8.0, '-8': 3.0, '-9': 11.0, '-10': 1.0, '-11': 12.0, '-12': 2.0, '-13': 1.0, '-20': 1.0} |
| KIN182 | 79 | 32 | | 100 | 0.79 | 0.017032191 | 0.57205453 | ACAAGGCAGCCTACATAATG | 58 | 58 | {'1': 2.0, '-1': 2.0, '-3': 31.0, '-5': 5.0, '-7': 9.0, '-8': 3.0, '-9': 9.0, '-10': 3.0, '-11': 8.0, '-12': 5.0, '-15': 2.0} |

| | |
|---|---|
| AVERAGE | 40.9 |
| STDEV | 7.279346735 |
| MEDIAN | 40.5 |

**Sanger Sequnecing results of sgRNA83822 and analysis on ICE**

| Label | ICE | KO-Score | KI-Score | ICE d | R Squared | Mean Discord Before | Mean Discord After | Guide Sequences | Control Sample Quality Score | Edit Sample Quality Score | Indels |
|---|---|---|---|---|---|---|---|---|---|---|---|
| KIN183 | 73 | 42 | | 100 | 0.73 | 0.010110835 | 0.588815177 | CAAGGCAGCCTACATAATGC | 58 | 58 | {'-1': 1.0, '-2': 6.0, '-3': 13.0, '-6': 3.0, '-7': 10.0, '-8': 8.0, '-9': 15.0, '-10': 6.0, '-11': 8.0, '-16': 2.0, '-19': 1.0} |
| KIN186 | 78 | 38 | | 100 | 0.78 | 0.004438409 | 0.656612301 | CAAGGCAGCCTACATAATGC | 58 | 61 | {'-2': 8.0, '-3': 14.0, '-6': 11.0, '-7': 8.0, '-8': 5.0, '-9': 15.0, '-10': 9.0, '-11': 7.0, '-13': 1.0} |
| KIN184 | 79 | 43 | | 100 | 0.79 | 0.012393583 | 0.627702943 | CAAGGCAGCCTACATAATGC | 58 | 61 | {'-1': 3.0, '-2': 6.0, '-3': 14.0, '-6': 6.0, '-7': 6.0, '-8': 1.0, '-9': 16.0, '-10': 12.0, '-11': 7.0, '-13': 2.0, '-16': 4.0, '-17': 2.0} |
| KIN185 | 79 | 46 | | 100 | 0.79 | 0.005756198 | 0.67214634 | CAAGGCAGCCTACATAATGC | 58 | 61 | {'-2': 10.0, '-3': 8.0, '-6': 10.0, '-7': 5.0, '-8': 6.0, '-9': 14.0, '-10': 7.0, '-11': 9.0, '-16': 9.0, '-18': 1.0} |
| KIN188 | 77 | 40 | | 100 | 0.77 | 0.008441895 | 0.616366727 | CAAGGCAGCCTACATAATGC | 58 | 58 | {'-1': 2.0, '-2': 11.0, '-3': 14.0, '-6': 5.0, '-7': 4.0, '-8': 1.0, '-9': 12.0, '-10': 12.0, '-11': 5.0, '-12': 2.0, '-13': 3.0, '-16': 2.0, '-18': 4.0} |
| KIN187 | 77 | 40 | | 100 | 0.77 | 0.014437428 | 0.613509282 | CAAGGCAGCCTACATAATGC | 58 | 58 | {'-2': 9.0, '-3': 19.0, '-6': 3.0, '-7': 10.0, '-8': 3.0, '-9': 11.0, '-10': 7.0, '-11': 6.0, '-12': 1.0, '-13': 2.0, '-16': 3.0, '-18': 3.0} |
| KIN191 | 79 | 48 | | 100 | 0.79 | 0.012107819 | 0.602215284 | CAAGGCAGCCTACATAATGC | 58 | 58 | {'-1': 2.0, '-2': 12.0, '-3': 14.0, '-6': 3.0, '-7': 8.0, '-8': 2.0, '-9': 10.0, '-10': 9.0, '-11': 7.0, '-12': 1.0, '-13': 1.0, '-16': 7.0, '-18': 3.0} |
| KIN190 | 79 | 31 | | 100 | 0.79 | 0.008510533 | 0.630216661 | CAAGGCAGCCTACATAATGC | 58 | 61 | {'-1': 1.0, '-2': 9.0, '-3': 8.0, '-6': 10.0, '-7': 2.0, '-8': 2.0, '-9': 30.0, '-10': 5.0, '-11': 11.0, '-13': 1.0} |
| KIN192 | 78 | 42 | | 100 | 0.78 | 0.008693021 | 0.664907087 | CAAGGCAGCCTACATAATGC | 58 | 61 | {'-2': 7.0, '-3': 16.0, '-6': 7.0, '-7': 13.0, '-8': 5.0, '-9': 10.0, '-10': 8.0, '-11': 7.0, '-12': 1.0, '-13': 2.0, '-18': 2.0} |
| KIN189 | 72 | 36 | | 100 | 0.72 | 0.008171226 | 0.638693207 | CAAGGCAGCCTACATAATGC | 58 | 61 | {'-1': 1.0, '-2': 6.0, '-3': 14.0, '-6': 4.0, '-7': 5.0, '-8': 4.0, '-9': 13.0, '-10': 8.0, '-11': 9.0, '-16': 3.0, '-18': 5.0} |

| | |
|---|---|
| AVERAGE | 40.6 |
| STDEV | 4.880801391 |
| MEDIAN | 41 |
