## Supplementary material for "Class III PI3K is essential for Wingless secretion and Evi/Wls recycling in *Drosophila*": Table S4

**Image parameters of STED pictures**

| **Figure #** | **3** | **4** | **5** | **5** | **6** | **6** | **6** |
|---|---|---|---|---|---|---|---|
| **Figure Panel** | **e-g** | **c-e** | **e-g** | **h** | **e-f (DMSO)** | **g-h (Dynasore2-24)** | **j-k (Bortezomib)** |
| Organism | *D. melanogaster* | *D. melanogaster* | *D. melanogaster* | *D. melanogaster* | *D. melanogaster* | *D. melanogaster* | *D. melanogaster* |
| Tissue | wing imaginal disc | wing imaginal disc | wing imaginal disc | wing imaginal disc | wing imaginal disc | wing imaginal disc | wing imaginal disc |
| Primary AB 1 | anti-Dlp ms | anti-wg ms 4D4, DSHB | anti-wg ms 4D4, DSHB | anti-wg ms 4D4, DSHB | anti-wg ms 4D4, DSHB | anti-wg ms 4D4, DSHB | anti-wg ms 4D4, DSHB |
| Secondary AB 1 | STAR635P ms | STAR635P ms | AF594 ms | AF594 ms | AF594 ms | AF594 ms | AF594 ms |
| Primary AB 2 | anti-GFP rb | anti-GFP rb | anti-GFP rb | anti-GFP rb | anti-GFP rb | anti-GFP rb | anti-GFP rb |
| Secondary AB 2 | AF594 rb | AF594 rb | STAR635P | STAR635P | STAR635P | STAR635P | STAR635P |
| Objective | 86x/1.2 HC PL APO CS2 (water) | 86x/1.2 HC PL APO CS2 (water) | 86x/1.2 HC PL APO CS2 (water) | 86x/1.2 HC PL APO CS2 (water) | 86x/1.2 HC PL APO CS2 (water) | 86x/1.2 HC PL APO CS2 (water) | 86x/1.2 HC PL APO CS2 (water) |
| Pixel Size | 0.0367x0.0367 micron^2 | 0.0367x0.0367 micron^2 | 0.0367x0.0367 micron^2 | 0.0367x0.0367 micron^2 | 0.0367x0.0367 micron^2 | 0.0367x0.0367 micron^2 | 0.0367x0.0367 micron^2 |
| Bits per pixel | 16-bit | 16-bit | 16-bit | 16-bit | 16-bit | 16-bit | 16-bit |
| XZ resolution | 27.2626 pixels per micron | 27.2626 pixels per micron | 27.2626 pixels per micron | 27.2626 pixels per micron | 27.2626 pixels per micron | 27.2626 pixels per micron | 27.2626 pixels per micron |
| Scan mode | xz | xz | xz | xz | xz | xz | xz |
| Zoom | 1.2 | 1.2 | 1.2 | 1.2 | 1.2 | 1.2 | 1.2 |
| Pixel dwell time | 2.5375 µs | 0.4625 µs | 0.4625 µs | 1.625 µs | 0.4625µs | 0.4625µs | 0.4625µs |
| Imaging mode | 3D STED | 3D STED | 3D STED | 3D STED | 3D STED | 3D STED | 3D STED |
| Line average | 1 | 1 | 1 | 1 | 1 | 1 | 1 |
| Line accumulation | 16 | 16 | 16 | 16 | 16 | 16 | 16 |
| Pinhole size | 64,8 µm | 70µm | 70µm | 70µm | 70µm | 70µm | 70µm |
| Excitation laser line | 580 nm / 638 nm | 580 nm / 638 nm | 580 nm / 638 nm | 580 nm / 638 nm | 580 nm / 638 nm | 580 nm / 638 nm | 580 nm / 638 nm |
| Excitation laser line intensity | 10% / 8% | 4% / 10% | 4% / 10% | 10% / 10% | 10% / 10% | 10% / 10% | 10% / 10% |
| Detector | HyD X2, 595-625 nm / HyD X4, 650-740nm | HyD X2, 595-625 nm / HyD X4, 650-740nm | HyD X2, 595-625 nm / HyD X4, 650-740nm | HyD X2, 595-625 nm / HyD X4, 650-740nm | HyD X2, 595-625 nm / HyD X4, 650-740nm | HyD X2, 595-625 nm / HyD X4, 650-740nm | HyD X2, 595-625 nm / HyD X4, 650-740nm |
| Tau gating excitation laser | 0.5-8ns | 0.5-8ns | 0.5-8ns | 0.5-8ns | 0.5-8ns | 0.5-8ns | 0.5-8ns |
| Depletion laser line | 775 nm | 775 nm | 775 nm | 775 nm | 775 nm | 775 nm | 775 nm |
| Depletion laser power | 65% | 75% | 75% | 75% | 75% | 75% | 75% |
| detection window | 3072x1536 pixel | 3072x2048 pixel | 3072x2048 pixel | 3072x2048 pixel | 3072x2048 pixel | 3072x2048 pixel | 3072x2048 pixel |
| Filter (Fiji) | gausion blur 1.5 | gausion blur 1.5 | gausion blur 1.5 | gausion blur 1.5 | gausion blur 1.5 | gausion blur 1.5 | gausion blur 1.5 |
